# Constriction forces imposed by basement membranes regulate developmental cell migration

**DOI:** 10.1101/2022.12.19.520983

**Authors:** Ester Molina López, Anna Kabanova, Maria D. Martín-Bermudo

**Affiliations:** Centro Andaluz de Biología del Desarrollo CSIC-Univ. Pablo de Olavide, Sevilla 41013, Spain; Department Physiology of Cognitive Processes, MPI for Biological Cybernetics, Tübingen, Germany

**Keywords:** laminins, migration, constricting forces

## Abstract

The basement membrane (BM) is a specialized extracellular matrix, which underlies or encase developing tissues. Mechanical properties of encasing BMs have been shown to profoundly influence the shaping of associated tissues. Here, we use the migration of the border cells (BCs) of the *Drosophila* egg chamber to unravel a new role of encasing BMs in developmental cell migration. BCs move between a group of cells, the nurse cells (NCs), that are enclosed by a monolayer of follicle cells (FCs), enveloped in turn by a BM, the follicle BM. We show that increasing or reducing the stiffness of the follicle BM, by altering laminins or Coll IV levels, conversely affects BC migration speed and alters migration mode and dynamics. Follicle BM stiffness also controls pairwise NC and FC cortical tension. We propose that constriction forces imposed by the follicle BM influence NC and FC cortical tension, which, in turn, regulate BC migration. Encasing BMs emerge as key players in the regulation of collective cell migration during morphogenesis.

## Introduction

The basement membrane (BM) is a dense, sheet-like form of an extracellular matrix (ECM) that underlies the basal side of epithelial and endothelial tissues and enwraps most developing organs (Hynes, 2012; Ozbek et al., 2010). BMs are highly conserved across evolution and are composed of a core set of proteins, including the secreted glycoproteins laminin, type IV collagen (Coll IV), entactin/nidogen and the heparan sulfate proteoglycan Perlecan (Yurchenco, 2011). Originally viewed as a static support ECM, BMs have recently been proven to act as active regulators of tissue shape and homeostasis.

Encasing BMs regulate the morphogenesis of the tissues they enwrap both molecularly, by modulating signaling to cells through surface receptors, and mechanically, by imposing patterned constricting forces. The best studied developmental function attributed to forces exerted by enwrapping BMs is the regulation of cell and organ shape (reviewed in (Sekiguchi and Yamada, 2018). Mechanically, BMs mediate organ shape by transmitting tension through interconnected cells. Involvement of BMs in organ shaping is well exemplified in the *Drosophila* central nervous system, wing and egg chamber development (Haigo and Bilder, 2011; Isabella and Horne-Badovinac, 2016; Meyer et al., 2014; Pastor-Pareja and Xu, 2011; Sanchez-Sanchez et al., 2017). Changes in the levels of several BM components, such as Collagen IV or Laminins, affect the shape of these tissues. Similarly, during mouse mammary gland duct development, BM accumulates at the base of duct buds, thereby constricting and elongating them (reviewed in (Muschler and Streuli, 2010). Finally, BMs also play critical roles in shaping biological tubes, including *Drosophila* embryonic salivary glands and renal tubes (Bunt et al., 2010; Urbano et al., 2009) and the *C. elegans* excretory system (Sundaram and Buechner, 2016). Besides changing in shape, cells of developing organs encased by BMs also undergo migratory processes. This is the case of the distal tip cell of the *C. elegans* somatic gonad (reviewed in (Sherwood and Plastino, 2018), the avian cranial neural crest cells (Hutchins and Bronner, 2019) or some cell populations during vertebrate branching morphogenesis (reviewed in (Wang et al., 2017). However, in contrast to the wealth of information of the role of BMs in cell migration when acting as a substratum, little is known about their function when acting as a “corset” encasing developing tissues.

The egg chamber (or follicle) of the *Drosophila* ovary, the structure that will give rise to an egg, has in recent years proven to be an excellent model system for investigating the *in vivo* roles of the forces exerted by encasing BMs in organogenesis (reviewed in (Montell, 2003). Egg chambers are multicellular structures that consist of 16-germline cell cysts, 15 nurse cells (NCs) and one oocyte, surrounded by a monolayer of somatic follicle cells (FCs), termed the follicular epithelium (FE). Each egg chamber progresses through fourteen developmental stages (S1-S14) (King, 1970). The apical side of FCs faces the germline, while the basal side contacts a BM produced by the FCs themselves, the follicle BM. The follicle BM of the *Drosophila* ovary contains Laminins, Coll IV, Perlecan and Nidogen (Gutzeit et al., 1991; Lerner et al., 2013; Lunstrum et al., 1988; Schneider et al., 2006). Initially, the egg chamber is spherical but between S5 and S10 elongates along its anterior-posterior (AP) axis, to acquire the final elliptical shape of the egg. During elongation, FCs move collectively over the inner surface of the follicle BM, a process that has been termed “global tissue rotation” (Haigo and Bilder, 2011). Rotation is accompanied by alignment of basal actin bundles and polarized secretion of BM components, which become organized in circumferential fibrils oriented perpendicular to the follicle’s AP axis (Cetera et al., 2014; Haigo and Bilder, 2011). The circumferential arrangement of fibrils is thought to act as a “molecular corset” that constrains follicle growth in the direction of the alignment, thus driving elongation. Once global tissue rotation has stopped and the circumferential fibrils of the corset are fully polarized and formed, around S9, a group of 6-10 anterior FCs, the border cells (BCs), delaminate from the epithelium and perform another type of directed collective cell migration (King, 1970; Montell et al., 1992). BCs extend actin-rich filopodia and migrate, as a group, in between the NCs, until they reach the oocyte. During BC migration, NCs exert compressive forces over the BCs influencing their migration (Aranjuez et al., 2016). Additionally, enzymatic removal of the follicle BM, using collagenase, has been shown to affect NC shape (Balaji et al., 2019). In this context, one could speculate that forces exerted by the follicle BM could in principle influence BC migration. However, this has never been tested.

Here, we show that increasing or reducing the stiffness of the follicle BM, by altering the levels of the BM components laminins or Coll IV, inversely affect BC migration speed. In addition, we found that follicle BM stiffness affects protrusion dynamics and migration mode. Furthermore, our results show that follicle BM stiffness controls pairwise FC and NC cortical tension. Finally, we show that a direct manipulation of FC tension also affects inversely BC migration speed. Based on these results, we propose that constriction forces imposed by the follicle BM affect FCs and NCs cortical tension, which, in turn, influences BC migration. Our results unravel a new role for the mechanical properties of the BMs enclosing developing organs, the regulation of the intercellular forces controlling collective cell migration. In addition, this work provides new insights towards our understanding of the remarkable and assorted ways whereby BMs regulate cell migration during morphogenesis.

## Results

### The stiffness of the follicle BM influences BC migration

To test the role of the forces exerted by the follicle BM on BC migration, we reduced the levels of two of its main components, Laminins or Coll IV. Laminins are large heterotrimeric glycoproteins that contain one copy each of an α, β and γ chain. *Drosophila* contains two different laminin trimers, each composed of one of the two α chains, encoded by the *wing blister* (*wb*) and *Laminin A* (*LanA*) genes, and the shared β and γ chains, encoded by the *Laminin B1 (LanB1)* and *Laminin B2 (LanB2)* genes, respectively (Chi and Hui, 1989; Graner et al., 1998; Kusche-Gullberg et al., 1992; Montell and Goodman, 1989). To reduce Laminin levels in follicles, we either expressed a LanB1 RNAi in all FCs by means of the *traffic jam* (*tj*) Gal4 line (*tj>LanB1^RNAi^*) or used a hypomorphic viable condition for the locus, LanB1^28a^/ l(2)K05404 (hereafter referred to as *LanB1^hyp^*), as both have been shown to drastically reduce LanB1 levels up to S8 (Diaz de la Loza et al., 2017; Urbano et al., 2009). Atomic force microscopy measurements have shown that follicle BM stiffness is significantly diminished in these mutant conditions from the germarium to S8 follicles (Diaz de la Loza et al., 2017). As here we found that LanB1 levels in the mutant follicles were kept constant and significantly inferior to controls until the end of oogenesis, S10 (Figure S1), we expected that the reduction in BM stiffness would also sustain until oogenesis is completed. To support our estimate of a reduction in BM stiffness in S9/S10 laminin-depleted follicles, we turned to an adapted organ swelling assay to assess overall BM stiffness (Crest et al., 2017). Immersion of live follicles in deionized water creates osmotic stress that leads to water influx into the follicle. Acute expansion of the organ challenges the BM, resulting in bursting which can be examined by live imaging (Crest et al., 2017). The frequency and speed at which the BM bursts has been hypothesized to reflect its overall stiffness. We found that S9-S10 LanB1-depleted follicles (*tj>LanB1^RNAi^* n=30 or *LanB1^hyp^* n=21, in this example and hereafter n indicates number of egg chambers analysed) burst more rapidly and frequently than wild type ones (n=38), strongly suggesting that BM stiffness was reduced in these mutant conditions at the time of BC migration (Figure S2A-C, F and G; Movie S1; (Crest et al., 2017).

Another key component of BMs is type IV collagen, a unique member of the collagen superfamily, which is composed of different α-chains that form distinct heterotrimers. The *Drosophila* Coll type IV trimer consists of two α1 chains and one α2 chain, encoded by the genes *Collagen at 25C* (*Cg25C*) and *Viking* (*Vkg*), respectively (Le Parco et al., 1986; Natzle et al., 1982; Yasothornsrikul et al., 1997). To decrease Coll IV levels, we expressed a Vkg RNAi in all FCs using the *tj*Gal4 driver (*tj>Coll IV^RNAi^*), an approach previously shown to efficiently reduce both Coll IV levels and BM stiffness (Crest et al., 2017; Haigo and Bilder, 2011). In fact, in agreement with previous analysis, we found that S9-S10 *tj>Coll IV^RNAi^* follicles burst more rapidly and frequently than wild type ones (n=18, Figure S2D, F and G; Movie S1; (Crest et al., 2017).

To quantify BC migration in different conditions in which the BM stiffness is altered, we performed live imaging of control and Laminin or Coll IV-depleted egg chambers. To visualize BCs, we generated transgenic flies that expressed the green fluorescent protein (GFP) driven by an enhancer of the *torso-like* (*tsl*) gene, which is active in BCs and in a group of posterior FCs (*tsl*GFP, see Materials and Methods, (Furriols et al., 2007). Quantification of BC migration speed from live imaging analysis revealed that BCs from *tsl*GFP; *tj>LanB1^RNAi^* (n=7), *tsl*GFP; *LanB1^hyp^* (n=6) or *tsl*GFP; *tj>Coll IV^RNAi^* (n=6) egg chambers moved faster than BCs from control follicles (*tsl*GFP; *tj>Gal4*, n=8) (Fig.1A–E, Movie S2, see Materials and Methods). In all experimental egg chambers, BCs completed their migration and reached the oocyte as in controls. Since we find that BC migration is similarly affected when reducing either laminin or Coll IV levels and as laminins are required for proper Coll IV deposition in the ovary (Diaz de la Loza et al., 2017), we continued our analysis using Laminin-depleted egg chambers.

**Figure 1.**
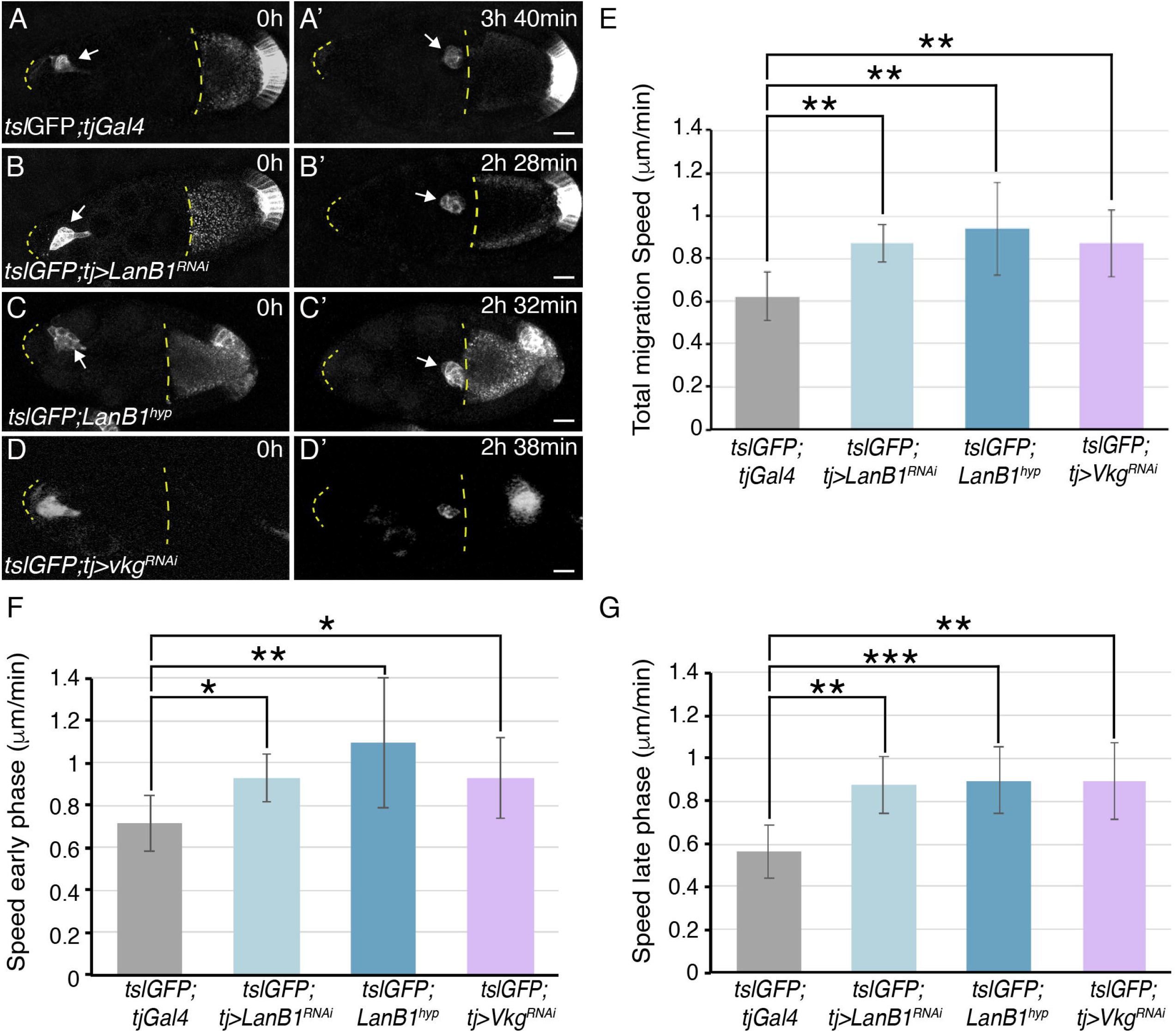
Reducing Laminins or Col IV levels results in increased BC migration speed. (A-D’) Stills taken from live imaging of migrating BCs from egg chambers of the indicated genotypes. (E, G) Quantification of BC migration speed during the whole migratory process (E) and during the early (F) and late (G) phases in egg chambers of the indicated genotypes. Scale bars in A’, B’, C’ and D’, 20μm.

BC migration to the oocyte can be divided in two phases. In an early phase-from detachment until halfway to the oocyte-movement is more streamlined and sliding. In the late phase-from midway to contacting the oocyte-movement is slower with clusters rotating in place (Bianco et al., 2007; Poukkula et al., 2011). We found that reducing laminin levels affected the migration speed of both phases (Fig.1F, G).

These results suggest that BM stiffness influences BC migration. However, the general follicle cell driver *tj-Gal4* directs expression in all FCs, including BCs. To rule out the possibility that the faster BC migration phenotype observed when depleting Laminins with *tjGal4* was due to a possible requirement of this BM component in BCs, we analysed the effects of expressing a *LanB1* RNAi specifically in BCs. To ensure early expression of the *LanB1* RNAi in BCs to achieve efficient RNAi knockdown in BCs, we combined the BC *Gal4* line *slbo-Gal4* (Montell et al., 1992) with two other *Gal4* lines, *C306Gal4i* and *torso-like-Gal4* (*tslGal4*), as they have been shown to be expressed early in oogenesis, before BCs detach from the follicular epithelium (Murphy and Montell, 1996; Laflamme et al., 2012; Aranjuez et al. 2012; Furriols et al. 2007). Live imaging showed that BCs from egg chambers expressing *LanB1^RNAi^* using this triple *Gal4* line combination (*triple>GFP; LanB1^RNAi^*, n=6) moved at similar speed than controls (*triple>GFP;* n=6, Figure S3, Movie S3). This result indicates that Laminins do not have an essential function in BCs, a finding that is consistent with the reported absence of Laminins expression in BCs during their migration (Medioni and Noselli, 2005).

To further test the idea that the stiffness of the BM affects BC migration, we analyzed the effect of increasing BM stiffness. To this end, we used the *tj-Gal4* to overexpress an mCherry tagged version of *EHBP1*(*EHBP1mCh*) in all FCs, an approach known to increase Coll IV fibril deposition and BM stiffness (Isabella and Horne-Badovinac, 2016, Crest, 2017 #1257). In fact, in agreement with those studies, we found that *tj>EHBP1mCh* follicles were more resistant to bursting than controls (Crest, 2017 #1257), Figure S2E-G, n=26). Live time-lapse imaging, showed that the overexpression of *EHBP1mCh* in all FCs delayed BC migration (*tslGFP; tj> EHBP1mCh*, Figure S4A-B’, Movie S4, n=6). However, as mentioned above, the general driver *tj-Gal4* drives expression in all FCs, including BCs (Figure S4). In this context, the BC migration defects observed in *tslGFP*; *tj>EHBP1mCh* follicles could also be due to an increase in Coll IV levels in BCs, rather than just a rise in follicle BM stiffness (Crest et al., 2017). To assess directly the impact of BM stiffness in BC migration, we used the *mirror-Gal4* (*mirr-Gal4*) driver, whose expression is restricted to a central region of the follicular epithelium (Borensztejn et al., 2013). We found that while BCs migrated normally anterior to the *mirror* region in the *tslGFP; mirr>EHBP1mCh* egg chambers (Fig.2A–C, n=77), migration speed was reduced once BCs reached the *mirror* region (Fig.2D, n=9, Movie S5). Furthermore, while migration was always completed in controls follicles (n=6), migration in the *mirror* region was halted in 20% of the experimental cases analysed (Fig.2E, n=9).

**Figure 2.**
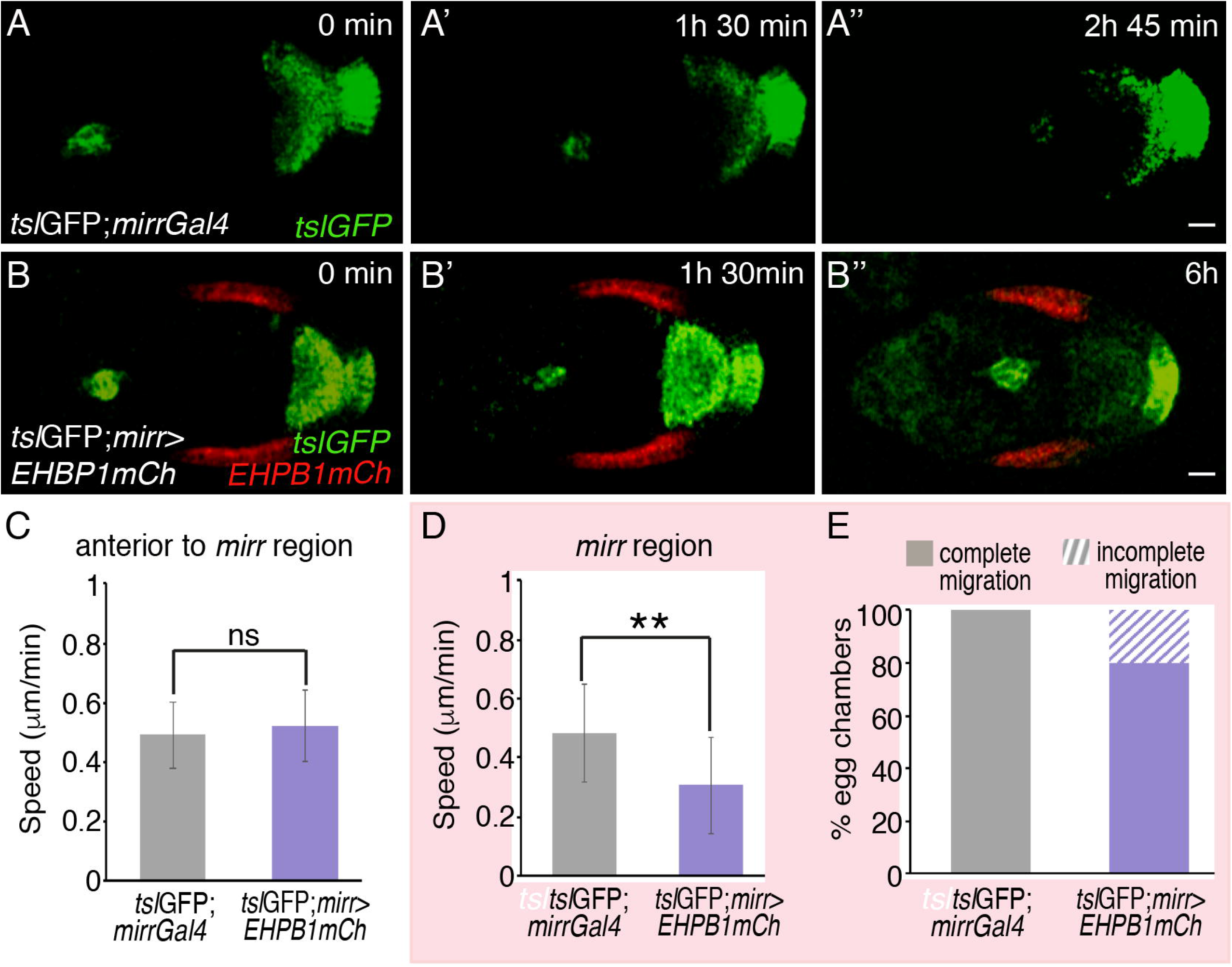
Increasing EHBP1 levels in FCs results in reduced BC migration. (A-B’’) Stills taken from live imaging of migrating BCs from egg chambers of the indicated genotypes. (C, D) Quantification of BC migration speed in egg chambers of the indicated genotypes in the area anterior to (C) and at the (D) *mirr* expressing region. (E) Quantification of egg chambers with complete BC migration. The statistical significance of differences was assessed with a t-test, * P value < 0.05. All error bars indicate s.d. Scale bars in A’’ and B’’, 20 μm.

### BM stiffness controls cellular protrusions dynamics and migration mode in BCs

Analysis of protrusions formation during the early and late phases of BC migration have shown an overall prominent bias in the direction of migration (Bianco et al., 2007; Prasad and Montell, 2007). As these protrusions extend between NCs, it is logical to think that any constrain(s) affecting NCs could influence the formation and dynamics of these protrusions. Thus, we next analyzed by live imaging the behavior of BC cellular protrusions in control and experimental clusters throughout the whole migratory process (Bianco et al., 2007; Fernandez-Espartero et al., 2013; Prasad and Montell, 2007). As shown previously, BC protrusions in control egg chambers showed an overall front bias (n=7, Fig.3A, E, Movie S2 (Bianco et al., 2007; Prasad and Montell, 2007). However, BCs from laminin-depleted egg chambers (*tsl*GFP; *tj>LanB1^RNAi^*) showed a significant increase in the number of protrusions extended laterally (n=7, Fig.3B, E). Moreover, BCs from *tslGFP; mirr>EHBP1mCh* egg chambers showed a decrease in the number of these lateral protrusions, only in the *mirror* expressing region (n=9) compared with controls (n=6) (Fig.3C, D, F).

**Figure 3.**
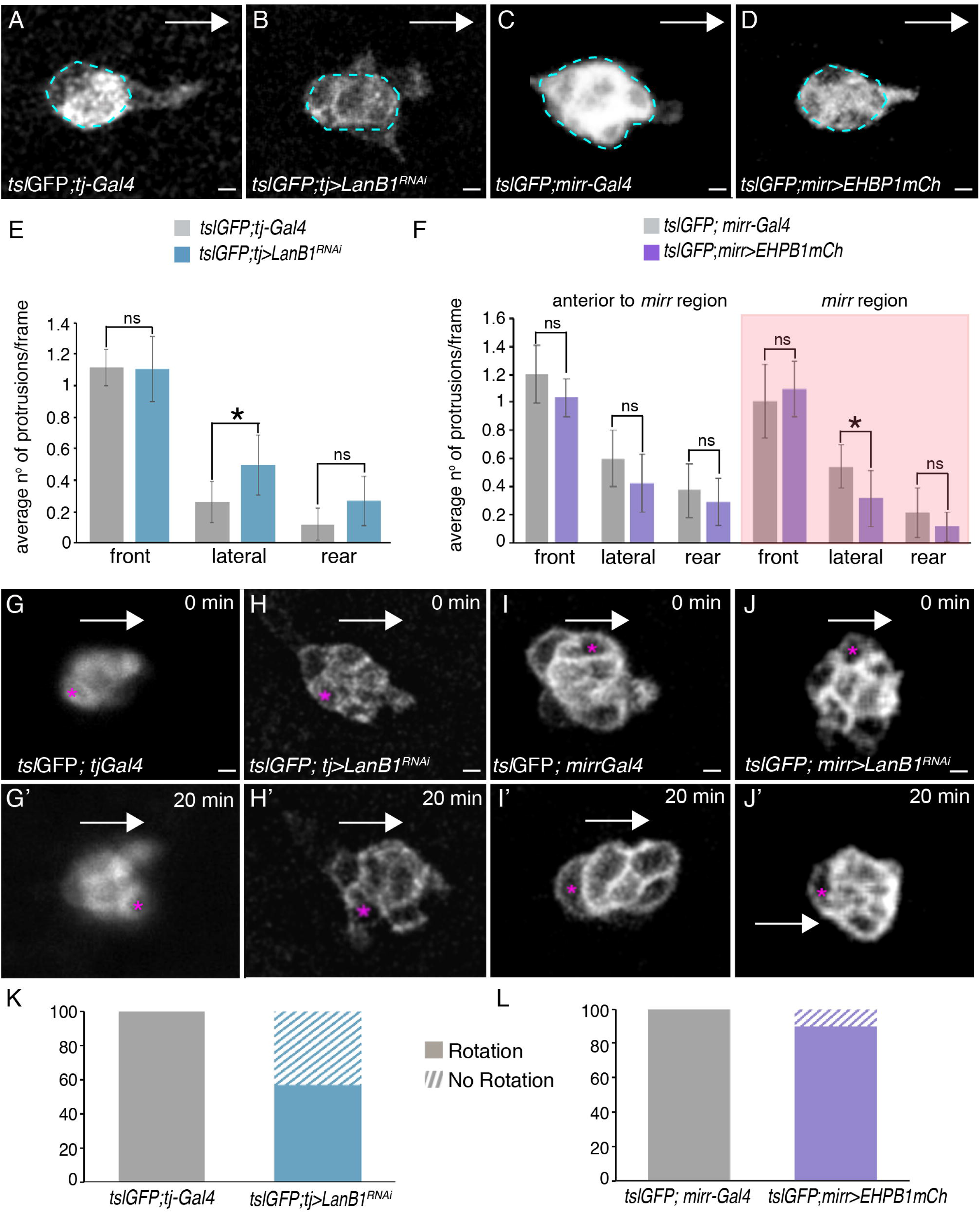
Modifying BM stiffness affects protrusion orientation and migration mode. (A-D) Stills taken from live imaging of BCs from egg chambers of the designated genotypes. (E-F) Mean number of protrusions observed per frame (snapshots) in each direction from BC clusters of egg chambers of the indicated genotypes. (G-J’) Stills taken from live imaging of BC clusters from egg chambers of the designated genotypes. (K, L) Quantification of BC cluster rotation. Scale bars in A-D and G-J, 10 μm.

As mentioned above, BCs show efficient forward movement during the early phase and increased tumbling during the late phase (Bianco et al., 2007). Next, we analyzed whether modifying the BM would affect the rotation mode. Using manual tracking (see Material and Methods), we found that, while control BC clusters always rotated (n=8, Fig.3G, G’, K, Movie S6), BC clusters in laminin-depleted egg chambers did not rotate in 42.8% of the cases analysed (n=8, Fig.3H, H’, K, Movie S6). In contrast, the slow-moving BC clusters of EHBP1 overexpressing follicles were still able to rotate (Fig.3I–J, L, n=10, Movie S7).

Altogether, these results demonstrate that the stiffness of the follicle BM influences dynamics and mode of BC migration.

### BM stiffness influences NC cortical tension

The current model proposes that NCs exert compressive forces over BCs that are perpendicular to the migration pathway and that mechanically influence their migration (Aranjuez et al., 2016). In this scenario, because NCs are externally constrained not only by FCs but also by the BM, the increase in migration speed we observe upon reduction of BM stiffness may arise from a drop in NC cortical tension. To test this possibility, we first analysed to what extent stiffness of the BM can influence NC cortical extension. In wild type egg chambers, BM stiffness over the NCs has been shown to gradually decrease from early S9 to S10 in the anterior region of the egg chamber, while a group of around 50 anterior FCs, the so-called stretched FCs (StFCs), progressively flatten in an anterior to posterior wave (Chlasta et al., 2017; Grammont, 2007; Kolahi et al., 2009). To test whether these changes in BM stiffness affect NC cortical tension, we performed laser ablation experiments in NCs under the stretching FCs throughout the flattening process (Figure S4A-F, (Farhadifar et al., 2007); see Materials and Methods, Movie S8). All cuts were made perpendicular to the AP axis (Figure S4) and the behaviour of cell membranes, visualised with Resille-GFP (Morin et al., 2001), was monitored up to 10 seconds after ablation (Movie S8). We found that, although some differences in BM stiffness were reported for mid and late S9 egg chambers, relative to early S9 (Chlasta et al., 2017), NC cortical tension did not change significantly throughout these stages, which is when BCs are migrating (n=16, Figure S5A-F). However, we did detect a significant decrease in NC cortical tension at S10 (n=15, Figure S5F), when anterior FCs are totally flattened, BC migration is completed and BM stiffness is much lower than that found at earlier stages (Chlasta et al., 2017). This result demonstrates that a reduction in BM stiffness below a certain threshold can influence NC cortical tension. Based on this result, we next tested whether the decrease in BM stiffness detected in the laminin-depleted egg chambers affected NC cortical tension. BM stiffness over the NCs has been found to be homogenous along the A/P axis in early S9 control egg chambers (Chlasta et al., 2017) (Lamire et al., 2020). In line with these results, we found that NC cortical tension was also homogenous along the A/P axis at this stage (n=14, Figure S5G). Thus, we decided to perform the next set of laser ablation experiments in NCs located in the central region of early S9 follicles, ahead of migrating BCs, as this will be the environment BCs will encounter during their movement towards the oocyte (Fig.4A). We found that the cortical tension of NCs in laminin-depleted follicles (Resille-GFP; *tj>LanB1^RNAi^*, n=17) was lower than in controls (Resille-GFP; *tj*, n= 15, Fig.4B–D, Movie S9). Accordingly, the levels of the phosphorylated form of the regulatory chain of nonmuscular Myosin II (encoded by the gene *spaghetti squash*, pSqh) at the NC cortex were lower in laminin-depleted follicles (n=10) compared with controls (n=9) (Figure S6B). An increase in NC contraction was shown to increase the levels of pSqh at the periphery of the BC cluster (Aranjuez et al., 2016). Accordingly, here we found that pSqh levels were reduced in the BCs of laminin-depleted egg chambers (n=8) compared with controls (n=9) (Figure S6D, E).

**Figure 4.**
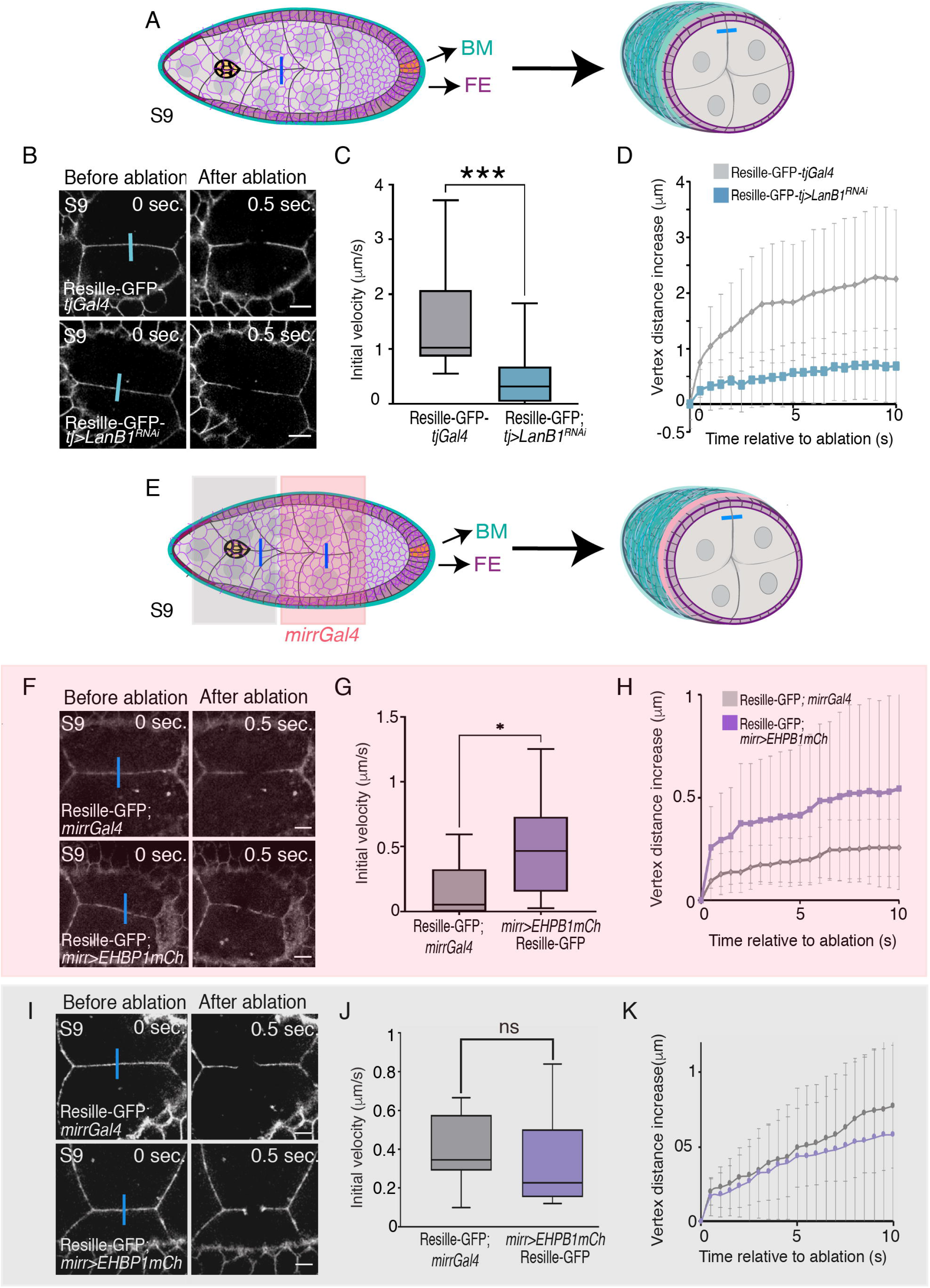
Changes in the stiffness of the BM affects NC cortical tension. (A) Schematic drawings of a S9 egg chamber illustrating the point of ablation (blue bar). NCs are in grey, BCs in yellow, FCs in purple and BM in blue. (B) Images of life S9 control (*tjGal4*) and *tj>LanB1^RNAi^* egg chambers expressing Resille-GFP before and after NC bonds are ablated. Blue bars indicate points of ablation. (C) Quantification of initial velocity of vertex displacement and (D) vertex displacement over time of the indicated ablated bonds. (E) Schematic drawing of a S9 egg chamber, similar to the one shown in A, illustrating the *mirror*Gal4 pattern of expression (*mirrGal4*, pink). (F, I) Life images of NCs located in the area anterior to (F) and at the (I) the *mirr* expressing region of S9 control (*mirr*Gal4) and *mirr>EHBP1mCh* egg chambers expressing Resille-GFP before and after NC bonds are ablated. Blue bars indicate points of ablation. (G, J) Quantification of initial velocity of vertex displacement and (H, K) vertex displacement over time of the indicated ablated bonds. The statistical significance of differences was assessed with a t-test, * P value < 0.05 and *** P value<0.001. All error bars indicate s.d. Scale bars in B F and I, 10 μm.

Altogether, our results support a model in which stiffness of the BM impacts on compressive forces exerted by NCs over the BCs, which is important for their movement. To further test this idea, we analyzed whether increasing BM stiffness could affect NC cortical tension. In order to do this, we expressed *EHBP1mCh* in the central region of the follicular epithelium and measured cortical tension and pSqh levels. Indeed, we found that the tension and pSqh levels at the cortex of NCs located under the *mirror* region in *mirr>EHBP1mCh* egg chambers (n=14 and n=7) was higher than that found in controls (n=13 and n=7, Fig.4F–H, Figure S6C), while no statistically significant difference was observed between experimental (n=13 and n=7) and controls (n=14 and n=7) NCs located anterior to the *mirror* region (Fig. 4I–K, Figure S6C). Furthermore, we found that the levels of pSqh were higher in BCs from *mirr>EHBP1mCh* (n=8, Figure S6F), being specifically significant at the lateral cluster periphery, where contraction forces from NCs might be higher in this experimental situation.

Finally, to further test our hypothesis that BM stiffness could influence BC migration via regulating NC cortical tension, we analysed whether a direct reduction of NC cortical tension would speed up BC migration. In order to do this, we expressed an RNAi against the Abelson interacting protein (Abi) in NCs, as its expression in FCs has been shown to cause a reduction in their cortical tension (Santa-Cruz Mateos et al., 2020). We found that the expression of an *Abi^RNAi^* RNAi in NCs (n=6, nos> *Abi^RNAi^* increased BC migration speed compared with controls (n=6) (Figure S7, Movie S10).

### BC migration defects found in laminin-depleted egg chambers are not due to changes in NC cytoplasmic pressure

Cell mechanical properties are not only dictated by cortical tension but also by cytoplasmic pressure. Collagenase treatment was shown to decrease NC cytoplasmic pressure, leading to suggest that BM stiffness could influence NC cytoplasmic pressure (Lamire et al., 2020). In this scenario, the effects on BC migration we observed when reducing laminins levels could also be due to alterations in NC cytoplasmic pressure. To test this possibility, we measured NC cytoplasmic pressure in early S9 control and *tj>LanB1^RNAi^* egg chambers. The radius of curvature of a spherical interface is inversely proportional to the difference in pressure between the two sides of the interface. Thus, to analyse NC cytoplasmic pressure, we determined the radius of curvature of the posterior membrane of NCs, by measuring-in 2D optical sections-the radius of the circle fitting this membrane, as described in (Lamire et al., 2020), Figure S8A, B, E, n>12 per position). In agreement with previous results (Lamire et al., 2020), we found a gradient of cytoplasmic pressure decreasing from anterior to posterior in early S9 control egg chambers (Figure S8E). This gradient was maintained in both laminin-depleted and *EHBP1mCh* overexpressing follicles (Figure S8A-F), which is consistent with previous results showing that altering the BM stiffness, by collagenase treatment, did not affect the gradient of NC cytoplasmic pressure (Lamire et al., 2020). However, even though absolute values of pressure were found lower in collagenase-treated follicles compared to untreated follicles (Lamire et al., 2020), we did not detect any significant difference between the curvature of NCs from *tj>LanB1^RNAi^* or *mirr>EHBP1mCh* follicles and that of controls (Figure S8E, F). To more directly address a possible role of NC cytoplasmic pressure on BC migration, we analysed BC migration in dicephalic (*dic^1^*) and kelch (*Kel^ED1^*) mutant egg chambers, as they have been shown to display reduced and increased NC cytoplasmic pressure, respectively (Lamire et al., 2020). Our *in vivo* analysis showed that the speed at which BCs moved in *dic^1^* and *Kel^ED1^* follicles was similar to that found in controls (Figure S8G-J, Movie S11). All together, these results suggest that NC cytoplasmic pressure on its own does not strongly contribute to the forces regulating BC migration.

### BM stiffness affects FCs’apical and basal cortical tension

As variations in BM mechanics have been shown to impact on FC shape (Chlasta et al., 2017), we wished to determine if follicle BM stiffness affected FC cortical tension. We first checked whether reducing BM stiffness altered basal FC tension by measuring cortical tension on the basal side of FCs in S9 control and laminin-depleted FCs. Laser cuts were made in main body cuboidal FCs contacting NCs just ahead of migrating BCs and perpendicular to the D/V axis (Fig.5A, A’). We observed that the tension at cell-cell contacts on the basal side of FCs in laminin-depleted egg chambers (*tj>LanB1^RNAi^*; Resille-GFP, n=15, Movie S12) was reduced compared to controls (*tj*; Resille-GFP, n=15, Fig.5B–D, Movie S12). These results demonstrated that follicle BM stiffness influences FC basal cortical tension. To further test this hypothesis, we assessed whether increasing BM stiffness could cause the opposite effect, a rise in basal FC tension. Indeed, we found that FCs expressing *EHBP1mCh* (Resille-GFP; *mirr>EHBP1mCh*, n=17) showed increased tension at basal FC-FC contacts compared to controls (Resille-GFP; *mirr*, n=16, Fig.5E–G).

**Figure 5.**
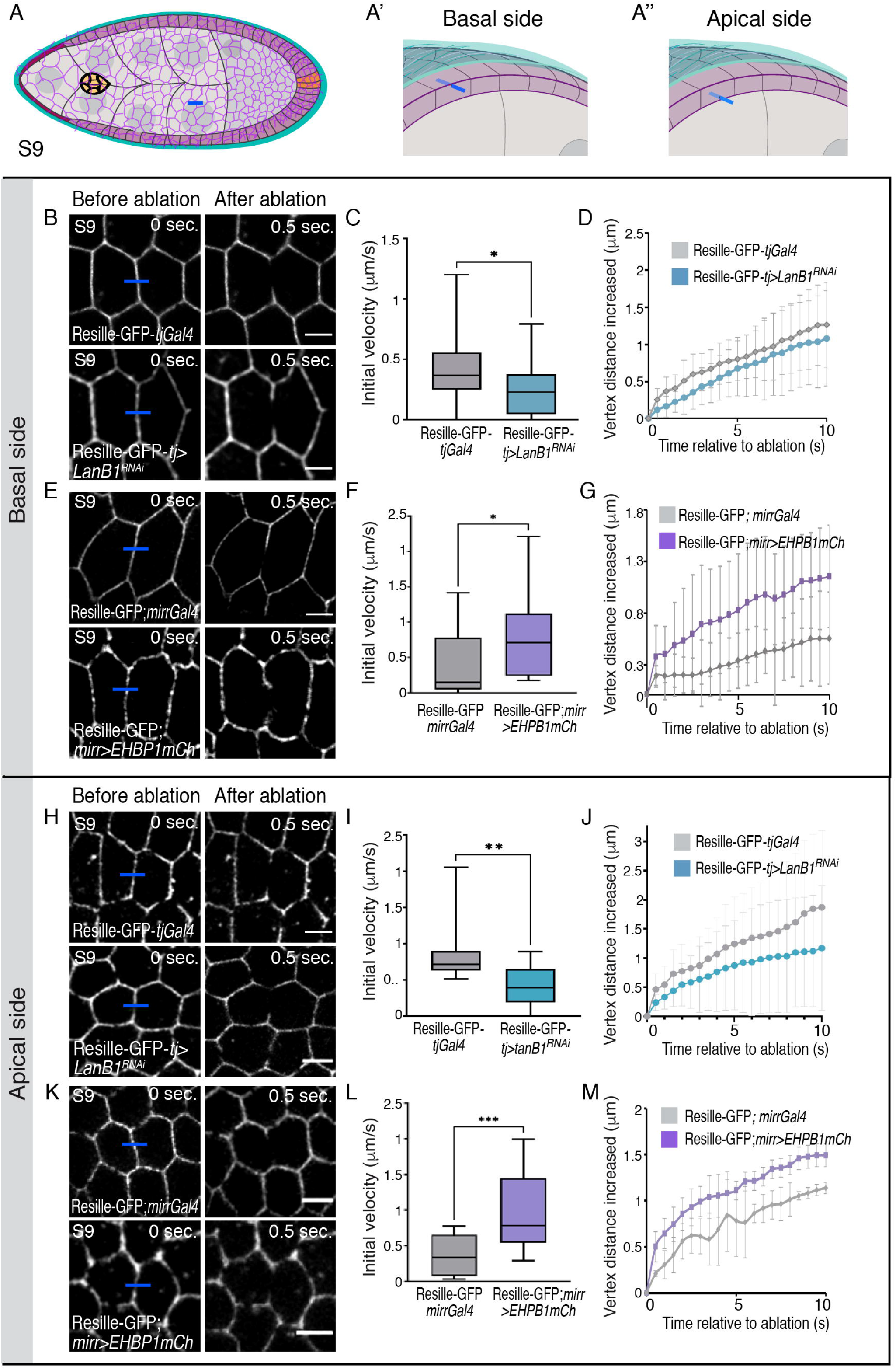
BM stiffness affects FCs membrane tension. (A-A’ ‘) Schematic drawings of a S9 egg chamber illustrating the point of ablation (blue bar) on the basal (A’) and apical (A’’) side of a FC. NCs are in grey, BCs in yellow, FCs in purple and BM in blue. (B, E, H, K) Images of the basal (B, E) and apical (H, K) sides of life S9 egg chambers of the indicated genotypes before and after FC bonds are ablated. Blue bars indicate points of ablation. (C, F, I, L) Quantification of initial velocity of vertex displacement and (D, G, J, M) vertex displacement over time of the indicated ablated bonds. The statistical significance of differences was assessed with a t-test, * P value < 0.05, ** P value < 0.001 and *** P value < 0.001. All error bars indicate s.d. Scale bars in B, E, H and K, 5μm.

We next tested whether manipulating BM stiffness could also affect tension at cell-cell contacts on the apical side of FCs (Movie S13). Recent studies have shown that decreasing Col IV levels reduced apical FC surface and caused defects in adherent junctions remodeling (Chlasta et al., 2017), suggesting a link between BM stiffness and FC apical tension. Similar to what happened on the basal side, our results showed that decreasing ((*tj>LanB1^RNAi^*; Resille-GFP, n=15) or increasing (*mirr>EHBP1mCh*, n=15) BM stiffness reduced or augmented respectively tension at apical FC-FC contacts (Fig.5H–M, Movie S13).

All together, these results suggest that the mechanical properties of the BM can also influence FC membrane tension. Because FCs are in direct contact with NCs, FC tissue mechanics could affect BC migration. To further test this hypothesis, we analyzed whether a direct manipulation of FC tension would affect BC migration. Expression of an *Abi* RNAi in all FCs leads to elimination of actin containing networks (Cetera et al., 2014; Squarr et al., 2016). Furthermore, we have recently shown that a reduction in *Abi* levels rescues the increase in FC cortical tension due to integrin elimination, suggesting that *Abi* may regulate cortical tension (Santa-Cruz Mateos et al., 2020). We thus analysed BC migration in follicles expressing an *Abi* RNAi in all FCs (*tslGFP*; *tj>AbiRNAi*). We found that BCs moved faster in *tslGFP*; *tj>AbiRNAi* follicles (n=8) compared to controls (*tslGFP*; *tj* Figure S9, Movie S14, n=9). While these results supported the idea that FC tension could influence BC migration, the fact that *tj-Gal4* drives gene expression in all FCs including BCs and that *Abi* depletion can block other collective migratory processes such as egg chamber rotation (Cetera et al., 2014), the defects observed in *tj>AbiRNAi* follicles could be, in part or totally, due to a specific role of *Abi* in BCs rather than an effect on FC tension. Thus, we used *mirr-Gal4* to deplete *Abi* only in central FCs (*mirr>Abi^RNAi^*). We first tested whether this approach altered FC tension of cell-cell contacts on the basal side of FCs. To this end, we performed laser ablation on the basal side of FCs located in the *mirror* area (Fig.6A). Indeed, we found that tension at FC-FC basal contacts was reduced in the Resille-GFP; *mirr>Abi^RNAi^* follicles (n=10) compared to controls (Resille-GFP; *mirr*, Fig.6B, C, n=13). Furthermore, laser ablation in NCs (Fig.6D) revealed that NC cortical tension was also reduced in the experimental egg chambers (Resille-GFP; *mirr>Abi^RNAi^*, n=15) compared to controls (Fig.6D–F, n=14). Finally, *in vivo* analysis of BC migration showed that BCs migrated faster in the *mirror* region (n=8) of experimental follicles compared to controls (n=8) (Fig.6G–I, Movie S15).

**Figure 6.**
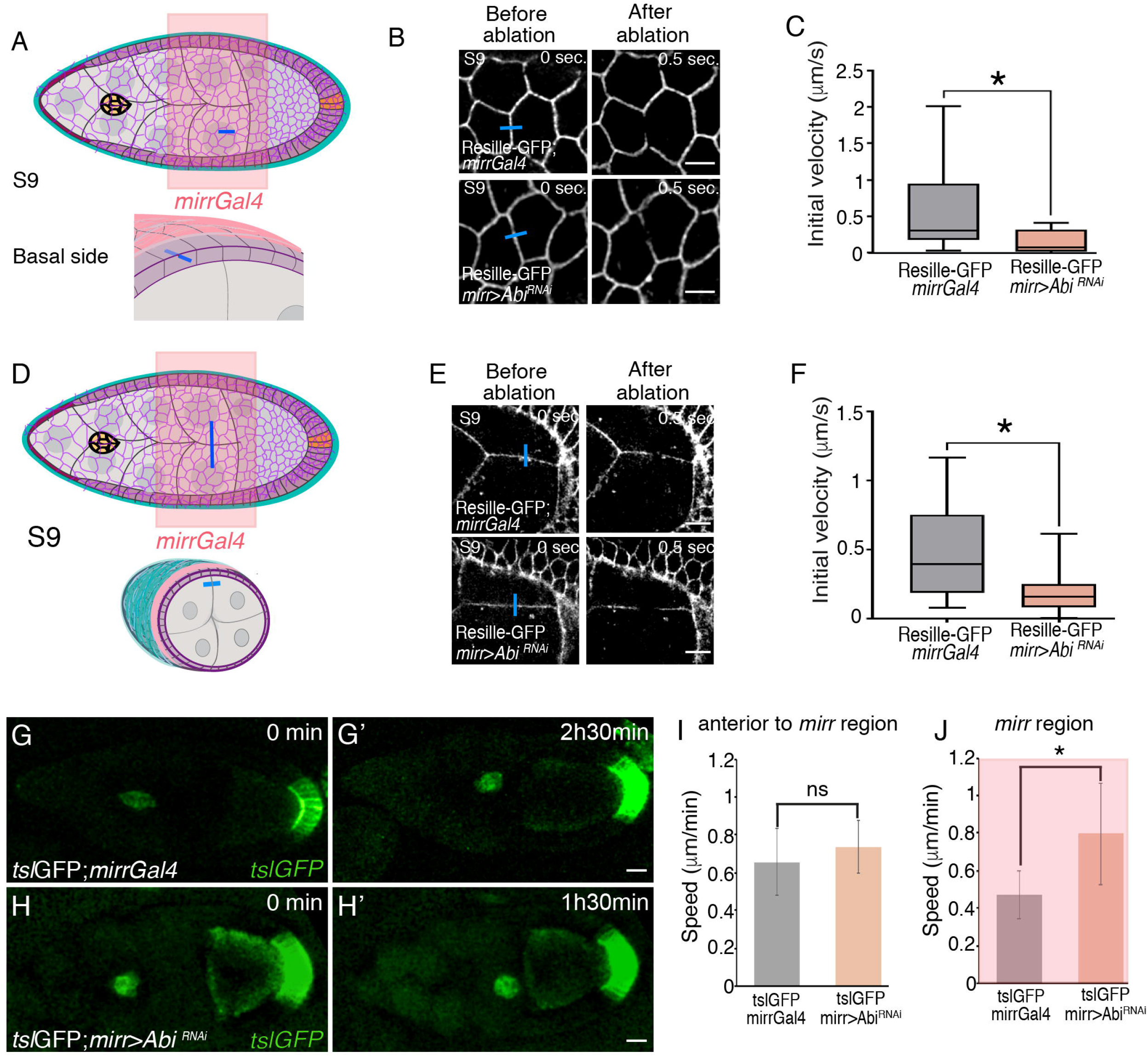
Reducing FCs membrane tension results in reduced NCs membrane tension and increased BC migration. (A) Schematic drawings of a S9 egg chamber illustrating the *mirror*Gal4 pattern of expression (*mirr*Gal4, pink) and the point of ablation in the basal side of FCs (blue bar). NCs are in grey, BCs in yellow, FCs in purple and BM in blue. (B) Images of life S9 control *mirr*Gal4 and *mirr>Abi^RNAi^* egg chambers expressing Resille-GFP before and after FCs bonds are ablated. Blue bars indicate points of ablation. (C) Quantification of initial velocity of vertex displacement of the indicated ablated bonds. (D) Schematic drawing of a S9 egg chamber illustrating the *mirrGal4* pattern of expression (pink) and the point of ablation in the NCs (blue bar). (E) Images of life S9 egg chambers of the indicated genotypes before and after NC bonds are ablated. Blue bars indicate points of ablation. (F) Quantification of initial velocity of vertex displacement of the indicated ablated bonds. (G-H’) Stills taken from live imaging of migrating BCs from egg chambers of the indicated genotypes. (I, J) Quantification of BC migration speed in the area anterior to (I) and at the (J) *mirr* expressing region. The statistical significance of differences was assessed with a t-test, * P value < 0.05, ** P value < 0.01 and *** P value < 0.001. All error bars indicate s.d. Scale bar in B, E and G-H’, 5μm, 10μm and 20μm, respectively.

## Discussion

BMs are thin, dense sheet-like types of ECM that underlie or surround virtually all animal tissues. By a combination of molecular and mechanical activities, BMs play multiple roles during tissue morphogenesis, maintenance and remodeling (Yurchenco, 2011). One of the most prominent developmental function attributed to BMs is cell migration. The conventional role of BMs during developmental migration is to provide an underlying substratum for moving cells. However, there are examples during morphogenesis where cells do not use BMs as a direct substrata, yet, the 3D environment through which they move is itself encased by a BM. Here, our results show that the stiffness of these encasing BMs can also influence collective cell migration, unravelling a new mechanical role for BMs during development.

The *Drosophila* egg chamber has served as a paradigm to study the role of encasing BMs in morphogenesis. Studies so far have revealed that the mechanical properties of the BM surrounding the egg chamber influence the behavior of cells in direct contact with it, the FCs. Constrictive forces applied by the egg chamber BM regulate changes in the shape of FCs that underlie tissue elongation (Haigo and Bilder, 2011; Isabella and Horne-Badovinac, 2016). BM stiffness has also been shown to affect the onset and speed of the collective movement of all FCs over the inner surface of the BM, during the process of “global tissue rotation” (Diaz de la Loza et al., 2017). Here, we show that the mechanical properties of the egg chamber BM can also regulate the behavior of cells that are not in direct contact with it, the BCs. BCs move using as a substratum NCs that are surrounded by FCs, encased, in turn, by a BM. Previous studies have shown that reciprocal mechanical interactions between the migrating BCs and the NCs substratum facilitate the movement of BCs (Aranjuez et al., 2016; Balaji et al., 2019). Our results reveal that constriction forces exerted by the BM also influence BC migration, as stiffness of the BM affects NC cortical tension. Recent studies have shown that BCs migrate along a central path where three or more NCs come together, as this is the path where the energy cost of unzipping NC-NC adhesion is most favorable (Dai et al. 2020). In this context, we could speculate that the reduction in NC cortical tension detected when reducing BM stiffness could weaken cell junctures, allowing for faster migration. Our results also show that even though NC cortical tension is reduced in laminin-depleted egg chambers, it seems to be maintained above the threshold necessary to preserve migration along the central path. We have also shown that affecting BM stiffness affect protrusion dynamics. A study analysing the requirements of Myo II in protrusion dynamics has suggested that, in contrast to the proposed role of protrusions as attachment sites regulating BC migration speed, protrusions function as sensory organs (Mishra et al., 2019). This work also implied that protrusions would extend in the direction of more advantageous environment, such as at the junctures where several NCs meet. In this scenario, as decreasing BM stiffness results in a reduction in NC cortical tension, this could explain our results showing a direct correlation between the number of protrusions oriented laterally and BM stiffness.

Besides cortical tension, cell mechanical properties are also dictated by cytoplasmic pressure. A gradient of cytoplasmic pressure within a defined group of NCs has been shown to induce TGFβ signaling in the surrounding epithelial cells. This, in turn, has been proposed to induce local softening of the BM and cell shape changes that promote elongation of the follicle, thus supporting a role for cytoplasmic pressure in shaping cells and organs (Lamire et al., 2020). However, here we show that cytoplasmic pressure on its own does not seem to compellingly influence BC migration. These results suggest that even though the different mechanical properties of cells are interconnected, each of them could impact the diverse cell biological processes in distinct ways. In fact, cytoplasmic pressure, and not cortical tension, has been proposed to regulate *Drosophila* nurse cell shape (Lamire et al., 2020). Similarly, during the process of mitotic rounding in the columnar epithelium of the *Drosophila* wing disc, cell area expansion was shown to be largely driven by cytoplasmic pressure, while roundness was primarily driven by cell-cell adhesion and cortical stiffness (Nematbakhsh et al., 2017). In the future, it will be interesting to understand the specific contribution of the different mechanical properties to the diverse morphogenetic processes.

Altogether, we would like to propose a model of the forces regulating BC migration in which, in addition to the already described forces that NCs apply directly over BCs and vice versa, more distant compressive forces exerted by the egg chamber BM emerge as another key regulator of this migratory process, by adjusting the mechanical properties of NCs and FCs (Fig.8). Our results prove that the 3D environment mechanically affecting developmental cell migration goes beyond adjacent cells or tissues, and suggest that forces exerted by all components of this complex 3D environment are most likely coordinated to allow proper cell migration.

We believe that the new mechanical role of the BM in cell migration proposed in this work is not just restricted to BCs, as, often, during morphogenesis, migrating cells move through tissues encased by a BM. During avian cranial neural crest migration, the BMs from the ectoderm and neural tube separate, yet, remain in close proximity forming a laminin-lined “channel” through which the cranial neural crest migrates (Hutchins and Bronner, 2019). Defective formation of this channel affects cranial neural crest migration (Hutchins and Bronner, 2019). We foresee that changes in the stiffness of the laminin-containing matrix lining the channel may also affect cranial neural crest migration. In the mouse cerebral cortex, BMs are found in the pia and around blood vessels (Sievers et al., 1994). Radial glia cells located near the ventricle extend long processes that attach to the pial BM. These cellular extensions serve as scaffold for the migration of Cajal-Retzius cells and neuroblasts, from the ventricular layer to the pial surface (Halfter et al., 2002). A major function of the pial BM is to provide an attachment site for the radial glia cell endfeet (Halfter et al., 2002). Damage to the pial BM results in defects in radial glial cell extension and consequently neuronal migration (Halfter et al., 2002). In this context, we believe that changes in the mechanical properties of the pial BM could also affect radial glial cell extension, which would result in neuronal migration defects.

The mechanical cues from the BMs have also been shown to contribute to the progression of several diseases including cancer. Epithelial cancer cells are bound within a BM and surrounded by a healthy stroma composed of collagen fibers and stromal cells. During the evolution of epithelial cancer invasion, cancer cells signal to the stromal cells to remodel the ECM at the tumor-stroma interface, loosen their cell-cell adhesion and breach the BM around the tumor and invade (reviewed in (Micalet et al., 2021). Experiments over the last years have focused on the impact of the stiffness of the ECM present at the tumor-stromal interface in tumor invasion. Thus, cell culture experiments have demonstrated that the rigidity of this ECM strongly affects the migration of glioma cells (Ulrich et al., 2009). Based on the results here presented, we would like to propose that the stiffness of the BM enclosing the tumor could also play an important role on the initiation of tumor invasion, and this will be worth to analyze in the near future.

Physical constraints imposed by BMs dictate many cellular events during tissue development. This is the case of branching morphogenesis in many mammalian organs where direct cell-matrix interactions, mainly mediated by integrins, orchestrate cellular rearrangements that sculpt the emerging tissue architecture (Nelson and Larsen, 2015). However, not all cells undergoing developmental cellular rearrangements are in direct contact with a BM. Instead, there are many examples in which a proportion of these cells contact other cells, which themselves are enclosed by a BM. This happens during the process of budding of the mammary gland, where the BM covers multilayered terminal ends growing dynamically by cellular rearrangements (Ewald et al., 2008). Similarly, in the stratified tip of pancreas and salivary glands, both outer, in contact with the BM, and inner tip cells rearrange and change position during the process of clefting (Hsu et al., 2013; Shih et al., 2016). Based on the results we present in this work, we foresee that the mechanical properties of the BM would also influence the rearrangement of cells located in the inner layers. Investigating this will help to better understand the complexity of morphogenesis in the context of multicellular developing tissues.

The results of this work unravel that, besides acting as a substratum, BMs can also regulate developmental cell migration by imposing constriction forces over the 3D environment through which cells move. A recent study, mining the Cancer Genome Atlas, has found that copy number alterations and mutations are more frequent in matrisome genes than in the rest of the genome, and that these mutations are statistically likely to have a functional impact (Izzi et al., 2020). Thus, identifying new roles for BMs in cell migration is helpful to understand not only morphogenesis but also cancer progression.

**Figure 7.**
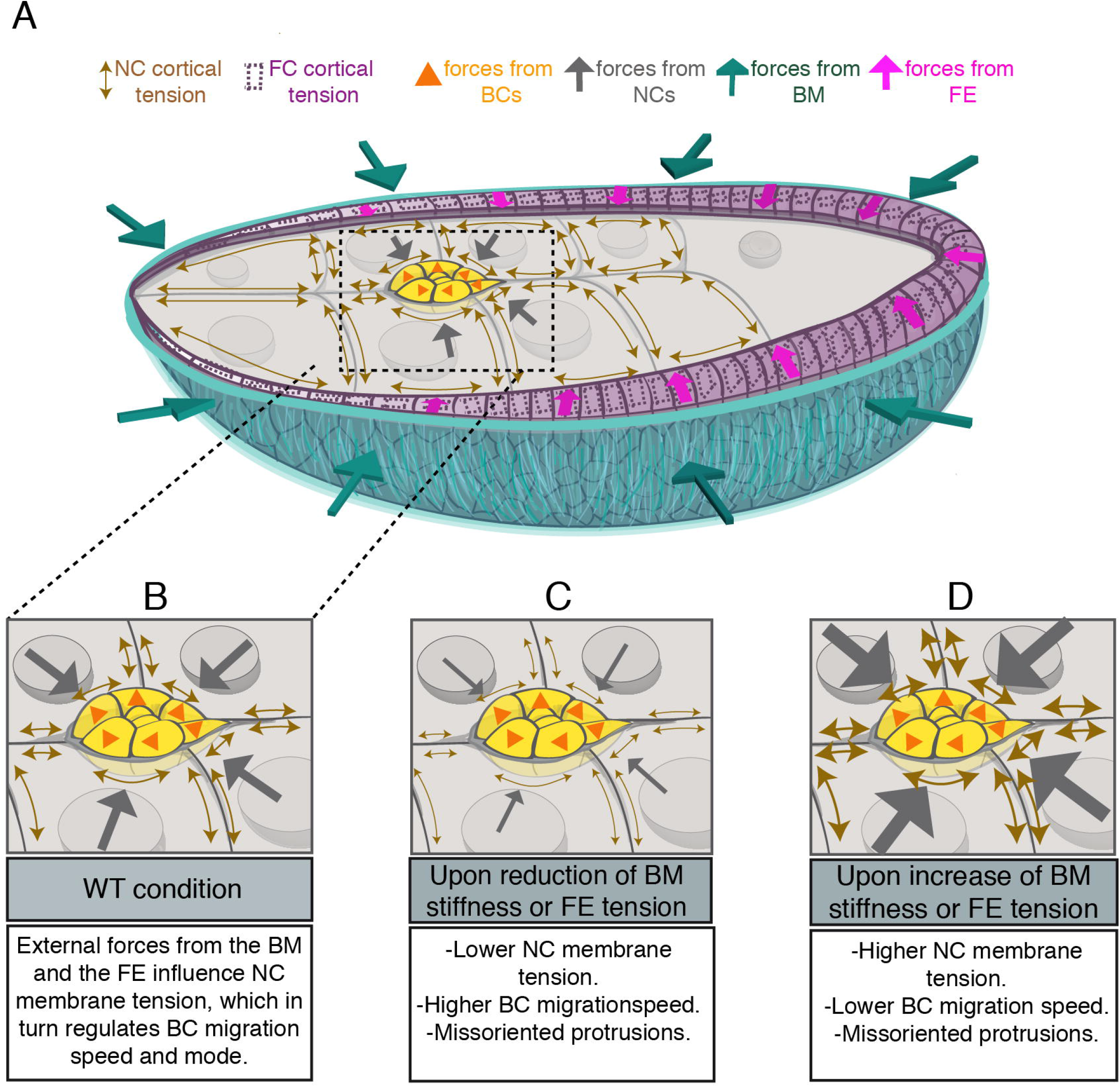
Proposed model of the forces exerted by the BM, FCs and NCs on BCs, and their effect on BC migration. (A) Proposed model for the forces exerted over BCs as they migrate. (B, C, D) Schematic drawing showing the consequences of altering the forces exerted over BCs on their migration.

## Acknowledgements

We acknowledge the Bloomington Stock Centre and the Developmental Studies Hybridoma Bank for fly stocks and antibodies. We also thank Sara Martín Villanueva and Marc Furriols for their help in generating the *tslGFP* construct and transgenic flies. Finally, we are grateful to A. González-Reyes for helpful remarks on the manuscript.

This work was supported by the Ministerio Español de Ciencia, Innovación y Universidades (MICINN, https://www.ciencia.gob.es/), grants numbers BFU2016-80797-R and PID2019-109013GB-100 to MDMB. EML was supported by a FPI studentship from the MICINN. We are also grateful to the MdM Unit of Excellence GEM (http://cellcollectives.com/) and to Junta de Andalucía (https://www.juntadeandalucia.es/) for their continuous funding and support.

## Author contributions

Conceptualization: M. D. Martín-Bermudo

Methodology: Ester Molina López, Anna Kabanova and M. D. Martín-Bermudo

Formal analysis: Ester Molina López and Anna Kabanova

Supervision: M. D. Martín-Bermudo

Writing - original draft, review & editing: M. D. Martín-Bermudo

Funding acquisition: M. D. Martín-Bermudo

## STAR Methods

### KEY RESOURCES TABLE

| REAGENT or RESOURCE | SOURCE | IDENTIFIER |
| --- | --- | --- |
| Antibodies |  |  |
| Rabbit anti-pSqh | Cell Signaling | 3671S |
| Anti-Rabbit Cy3 | Jackson ImmunoResearch Laboratories, Inc | 211-502-171 |
| Anti-LanB1 | Abcam | EPR3189 |
| Chemicals, peptides, and recombinant proteins |  |  |
| Rhodamine-Phalloidin | Molecular Probes | R37112 |
| Hoechst | Thermo Fisher | H21491 |
| Experimental models: Organisms/strains |  |  |
| *LanB1*<sup>28a</sup> (II) | (Urbano et al., 2009) | N/A |
| *l(2)K05404* | Bloomington Drosophila Stock Center | BDSC 13957 |
| *Kelch*<sup>ED1</sup> | Bloomington Drosophila Stock Center | 4893 |
| *dic*<sup>1</sup> | Bloomington Drosophila Stock Center | 4223 |
| *UAS-LanB1 RNAi* | VDRC | V23119 |
| *UAS-Vkg RNAi* | Bloomington Drosophila Stock Center | BDSC 50895 |
| *UAS-Abi RNAi* | DGRC Kyoto | 9749R |
| *UAS-EHBP1mCh* | Bloomington Drosophila Stock Center | BDSC 67145 |
| *mirror Gal4* | Bloomington Drosophila Stock Center | BDSC 63048 |
| *slbo Gal4* | Bloomington Drosophila Stock Center | BDSC 8915 |
| *C306 Gal4* | Bloomington Drosophila Stock Center | BDSC 3743 |
| *Tsl Gal4* | (Furriols et al. 2007) | N/A |
| *nos Gal4* | Bloomington Drosophila Stock Center | BDSC 4937 |
| ResilleGFP | Kindly provided by Prof. Wieschaus | N/A |

### RESOURCE AVAILABILITY

#### Lead Contact

Detailed information and requests for resources generated in this study should be addressed to and will be fulfilled by the lead contact María D Martín Bermudo.

#### Materials availability

All reagents generated in this study are available upon request to the lead contact.

#### Data and code availability

All data reported in this paper and any additional information required to reanalyze the data described in this paper is available from the lead contact upon request.

This paper does not report original code.

### EXPERIMENTAL MODEL AND SUBJECT DETAILS

#### *Drosophila* husbandry

*Drosophila melanogaster* strains were raised at 25 °C on standard medium. Laminin hypomorphs were trans-heterozygous for lanB1^28a^ (Urbano et al., 2009) and l(2)k05404 (Spradling et al., 1999), neither of which affect the coding region. To knock down laminin, Coll IV or Abi levels or to increase EHBP1 expression in the follicular epithelium, the *traffic jam* (*tj*) (Li et al., 2003) and the *mirror* (*mirr*) (Borensztejn et al., 2013) Gal4 drivers were used in combination with the following lines: *UAS-LanB1 RNAi* (VDRC 23119), *UAS-Vkg RNAi* (BDSC 50895), *UAS-abi RNAi* (DGRC-Kyoto 9749R) and *UAS-EHBP1mch* (BDSC 67145). To visualize cell membranes in vivo, using the membrane marker Resille-GFP, and, at the same time, use the *tj* Gal4 driver to express a desired UAS construct in all FCs, we generated flies with a recombinant 2^nd^ chromosome carrying both transgenes, Resille-GFP-*tj*. Stocks used in this study are described in the table.

#### Generation of *Drosophila* transgenic lines expressing GFP in border cells

To generate the *tslGFP* construct, an EcoR1/BamH1 fragment from C4PLZ-*tsl*(G), previously shown to drive expression in BCs (Furriols et al., 2007), was subcloned into the pCasper-GFP-PH vector (Barolo et al., 2004). The plasmid was introduced into the germ line of *w^1118^* flies by the company BestGene Inc following standard methods and several independent transgenic lines were isolated.

### METHODS DETAILS

#### Immunohistochemistry

Flies were grown at 25°C and yeasted for 2 days prior to dissection. Ovaries were dissected at room temperature in Schneider’s medium (Sigma Aldrich). After that, fixation was performed incubating egg chambers for 20 minutes in 4% paraformaldehyde in PBS (ChemCruz). Samples were then permeabilized with a phosphate-buffered saline +1% Tween20 solution (PBT) and incubated in a blocking solution (PBT10) for 1 hour. Ovarioles were then incubated with the primary antibody rabbit anti-phosphomyosin (Cell Signalling) overnight and, following a washing step, a secondary antibody Cy3 (Jackson ImmunoReseach Laboratories, Inc.) was added. To visualize F-acting and DNA, ovarioles were incubated for 30 minutes with Rhodamine Phalloidin (Molecular Probes, 1:40) and 10 minutes with Hoechst (Molecular Probes, 1:100), respectively. Experimental and control ovarioles were always mixed and stained together. Samples were mounted in Vectashield (Vector Laboratories) and imaged on a Leica SPE confocal microscope (DM 2500).

#### Time-lapse imaging

For time-lapse imaging, 1-2 days old flies were fattened on yeast for 24-48 hours before dissection. Ovaries were dissected in Schneider’s medium. Culture conditions were performed as described in (Prasad and Montell, 2007). Movies were acquired on a Leica SP5 MP-AOBS confocal microscope with a 40 × 1, 3 PL APO objective and a Nikon confocal microscope with a 40x oil objective. Z stacks of 30-40 slices (0.75-1.5 μm interval) were taken every 3-4 minutes.

#### Osmotic swelling assay

For the osmotic swelling assays, flies were prepared and dissected as described in the Time-lapse imaging section. After dissection, ovarioles were mounted in a Petri Dish and the Schneider’s medium was quickly replaced with dH_2_O (Crest et al., 2017). Images were collected every 30 seconds during 1 hour on a Zeiss Axioimager fluorescence microscope with an 5X air objective.

#### Laser ablation

Laser ablation experiments were performed in an Olympus IX-81 inverted microscope equipped with a spinning disk confocal unit (Yokogawa CSU-X1), a 100x oil objective, a 355 nm pulsed laser, third-harmonic, solid-state UV laser and a Evolve 512 EMCCD digital camera (Photometrics). To analyze tension in NCs and FCs, a pulse of 1000 μJ energy and 20 msec. and 75 μJ energy and 4 msec. duration, respectively, was applied to sever plasma membranes of cells. In all cases, cell surfaces were visualized with the membrane marker Resille-GFP and a Cobolt Calypso state laser (l= 491 nm 50 mW) was used for excitation of the GFP. To minimize potential effects due to anisotropic distribution of forces in NCs and FCs, cuts were made perpendicular to the anterior-posterior axis and to the dorsal ventral axis of the egg chamber, respectively. Images were taken 3 sec before and 10 sec after laser pulse, every 0,5 seconds. To analyze the vertex displacements of ablated cell bonds, the vertex distance increase from different ablation experiments (DL) was averaged using as L_0_ the average of distance of the vertexes 3 sec before ablation. The initial velocity was estimated as the velocity at the first time point (t1=0,5 sec). Standard deviation (s.d.) was determined.

#### Image processing and data analysis

Measurements of cluster velocity, cluster rotation and protrusion dynamics were done manually as described in (Fernandez-Espartero et al., 2013). Briefly, once the body of the cluster was defined according to (Poukkula et al., 2011), forward directed speed was calculated using the distance between the centre of the cluster at one time point and the next in the X axis only. When needed, drift of egg chambers was corrected using the plugin StackReg from FIJI-Image J (Thevenaz et al., 1998). To study BC cluster rotation, the position of an individual BC was manually tracked in z stacks of GFP confocal sections time point by time point for an interval of 20 minutes during the late phase of migration. To analyze protrusion orientation, a line was drawn from the centre of the cluster to the tip of the extension. The angle of this line relative to the x-axis was calculated. Extensions were classified as front (45°-315°), side (255°-315° and 45°-135°) and back (135°-225°). In all cases, an average of 150 protrusions was analyzed.

Analysis of the radius of curvature of NCs was carried out by measuring the radius of manually drawn circles that fit the curvature of the posterior facing membranes of NCs. This quantification was done only on one NC per row (anterior, central anterior, central posterior and posterior) in at least 12 independent follicles.

For the quantification of pSqh fluorescent intensities in NCs, square regions of interest (ROIs) of 28.94 μm2 (20×20 pixels) were applied to the membranes of central-posterior and posterior NCs. The mean grey value was then measured using the FIJI-Image J measure tool. This analysis was performed in individual Z-confocal sections corresponding to focal planes in which BCs were visible. For the quantification of pSqh fluorescence intensity in BCs, maximum projections of 5 Z-confocal sections were taken every 0.75 μm. Then 28.94 μm2 square ROIs were applied to the front, rear, and both sides of BCs and the mean grey value was then measured with the FIJI-Image J measure tool.

Statistical analysis of significant differences between control and experimental samples was done using Student’s *t*-test.

## Supplemental Information

**Movie S1 Osmotic-swelling experiments**

Osmotic bursting tests of egg chambers of the designated genotypes immersed in deionized water, related to Figure S1.

**Movie S2 *In vivo* BC migration in control and laminin- and Col IV-depleted egg chambers**

BC migration in control (*tsl*GFP; *tjGal4*) and laminin (*tsl*GFP; *tj>LanB1^RNAi^*, *tsl*GFP; *Lanb1^hyp^* or Col IV (*tsl*GFP;*tj>Col IV RNAi*) depleted egg chambers, related to Figure 1.

**Movie S3 *In vivo* migration of control and *LanB1^RNAi^* expressing BCs**

Migration of control (*C306; slbo; tsl;GFP*) and *LanB1^RNAi^* expressing BCs (*C306;slbo;tsl;GFP>LanB1 RNAi*), related to Figure S2.

**Movie S4 *In vivo* BC migration in control and *tj>EHBP1mCh* egg chambers**

BC migration in *tsl*GFP; *tjGal4* and *tsl*GFP; *tj>EHBP1mCh* egg chambers, related to Figure S3. *tsl*GFP is in green and *EHBP1mCh* in red, related to Figure S3.

**Movie S5 *In vivo* BC migration in control and *mirr>EHBP1mCh* egg chambers**

BC migration in *tsl*GFP; *mirrGal4* and *tsl*GFP; *mirr>EHBP1mCh* egg chambers, related to Figure 2. *tsl*GFP is in green and *EHBP1mCh* in red, related to Figure 2.

**Movie S6 BC cluster rotation in control and laminin-depleted egg chambers**

BC cluster rotation in control (*tsl*GFP; *tjGal4)* and laminin depleted (*tsl*GFP; *tj>LanB1RNAi)* egg chambers, related to Figure 3.

**Movie S7 BC cluster rotation in control and EHBP1mCh overexpressing**

BC cluster rotation in control (*tsl*GFP; *mirrGal4*) and *EHBP1mCh* overexpressing (*tsl*GFP; *mirr>EHBP1mCh*) egg chambers, related to Figure 3.

**Movie S8 Laser ablation of cell bonds between NCs of control egg chambers.**

Movies correspond to the ablation experiment shown in Figure S4. NCs membranes are visualised with Resille-GFP. A cell bond between two control NCs is ablated. GFP fluorescent is lost in the middle of the ablated bond upon laser ablation. The movie continues 10s after the cut and shows displacement of the vertexes. Images are taken every 0.5 seconds.

**Movie S9 Laser ablation of cell bonds between NCs of control and laminin-depleted egg chambers.**

Movies correspond to the ablation experiment shown in Figure 4. NCs membranes are visualised with Resille-GFP. A cell bond between two control NCs is ablated. Upon laser ablation, GFP fluorescent is lost in the middle of the ablated bond. The movie continues 10s after the cut and shows displacement of the vertexes. Images are taken every 0.5 seconds.

**Movie S10 *In vivo* BC migration in control and *nos>abiRNAi* chambers**

BC migration in *nos;hisYFP* and *nos>abiRNAi;hisYFP* egg chambers, related to Figure S6.

**Movie S11 *In vivo* BC migration in control and in *dic^1^* and *Kel^ED1^* mutant egg chambers**

BCs were directly visualized using bright field, related to Figure S7.

**Movie S12 Laser ablation on the basal side of cell bonds between FCs of control and laminin-depleted egg chambers.**

Movies correspond to the ablation experiment shown in Figure 5. The membranes on the basal side of FCs are visualised with Resille-GFP. A cell bond between two control FCs is ablated. GFP fluorescent is lost in the middle of the ablated bond upon laser ablation. The movie continues 10s after the cut and shows displacement of the vertexes. Images are taken every 0.5 seconds.

**Movie S13 Laser ablation on the apical side of cell bonds between FCs of control and laminin-depleted egg chambers.**

Movies correspond to the ablation experiment shown in Figure 5. The membranes on the apical side of FCs are visualised with Resille-GFP. A cell bond between two control FCs is ablated. GFP fluorescent is lost in the middle of the ablated bond upon laser ablation. The movie continues 10s after the cut and shows displacement of the vertexes. Movie length and frame rate are as described for Movie S12.

**Movie S14 *In vivo* BC migration in control and *tj>abiRNAi* chambers**

BC migration in *tsl*GFP; *tjGal4* and *tsl*GFP; *tj>abiRNAi* egg chambers, related to Figure S8.

**Movie S15 *In vivo* BC migration in control and *mirr>abiRNAi* chambers**

BC migration in *tsl*GFP; *mirrGal4* and *tsl*GFP; *mirr> abiRNAi* egg chambers, related to Figure 6.

**Figure S1. Quantification of laminin levels in S9/S10 *laminin-depleted* ovaries.**

(A) S9/S10 control *tjGal4* and (B) *tj>LanB1RNAi* (B) egg chambers stained with anti-LanB1 antibody (green), the DNA marker hoechst (blue) and the F-actin marker Rhodamine-Phalloidin (F-actin, red). (C) Quantification of the LanB1 levels in egg chambers of the specified genotypes. The statistical significance of differences was assessed with a t-test, **** P value<0.0001. Scale bars in A and B, 20μm.

**Figure S2. Reducing laminin levels in FCS results in a decrease in BM stiffness.**

(A-E) Stills taken from live imaging of osmotic bursting of follicles of the designated genotypes when placed in deionized water. (F, G) Quantification of the percentage (F) and frequency (G) of bursting. Scale bars in A-E, 50μm.

**Figure S3. Decreasing laminin levels in BCs does not affect their migration.**

(A-B’’) Stills taken from live imaging of BCs from control (*C306Gal4; slboGal4; tslGal4*) and *C306; slbo; tsl>LanB1^RNAi^* egg chambers. (C) Quantification of the migration defects in egg chambers of the indicated genotypes. Scale bars in A’’ and B’’, 20μm.

**Figure S4. Elevating EHBP1 levels in all FCs results in reduced BC migration speed.**

(A-B’) Stills taken from live imaging of migrating BCs from egg chambers of the indicated genotypes. Scale bar in A’ and B’, 20μm.

**Figure S5. NC cortical tension throughout egg chamber development from early S9 to S10.**

(A-D) Schematic drawings of early S9 (A), middle S9 (B), late S9 (C) and S10 egg chamber illustrating the BCs (yellow), NCs (grey), FCs (purple), BM (green) and the point of ablation in the NCs (blue bar). (A’-D’) Images of life control egg chambers of the indicated developmental stages expressing Resille-GFP before and after NC bonds are ablated. Blue bars indicate points of ablation. (E) Schematic representation of an egg chamber and the point of ablation in NCs. (F, G) Quantification of the initial velocity of vertex displacement of the indicated ablated bonds. Scale bar in A’-D’, 10μm.

**Figure S6. Modifying BM stiffness affects pSqh levels at the cortex of NCs**

(A, D) S9 control *tjGal4* egg chambers stained with anti-pSqh (red) and the DNA marker hoechst (blue). (B, C, E, F) Quantification of pSqh levels in the NCs (B, C) and in the BCs (E, F) of egg chambers of the specified genotypes. Scale bar in A, 20μm.

**Figure S7. Decreasing *abi* RNA levels in NCs results in increased BC migration speed.**

(A-B’) Stills taken from live imaging of migrating BCs from egg chambers of the indicated genotypes. (C) Quantification of the migration defects in egg chambers of the specified genotypes. Scale bars in A’ and B’, 20μm.

**Figure S8. Contribution of NC cytoplasmic pressure to BC migration**

(A-D) S9 controls *tjGal4*, *tj>LanB1^RNAi^*, *mirrGal4* and and *mirr>EHBP1mCh* egg chambers stained with the F-actin marker Rhodamine-Phalloidin (F-actin, red) and the DNA marker hoechst (blue). Arrows in A indicate the curvature of the membranes between an anterior (A) and an anterior central NC (Ca); between two central NCs-a Ca and a Cp (a posterior central NC); between a Cp and a posterior (P) NC and between a P NC and the oocyte membrane. (E, F) Quantification of the radius of curvature of A, Ca, Cp and P NCs in S9 egg chambers of the indicated genotypes. BC clusters are marked with a red circle. (G-I) Stills taken from live imaging of migrating BCs from egg chambers of the indicated genotypes. (J) Quantification of BC migration speed in egg chambers of the indicated genotypes. Scale bars in A-D and G-I’, 20 μm.

**Figure S9. Decreasing *abi* RNA levels in all FCs results in increased BC migration speed.**

(A-B’) Stills taken from live imaging of migrating BCs from egg chambers of the indicated genotypes. (C) Quantification of the migration defects in egg chambers of the specified genotypes. Scale bars in A’ and B’, 20μm.

