## Supplementary figures and images for "Constriction forces imposed by basement membranes regulate developmental cell migration"

### Sup. Fig.1.tif

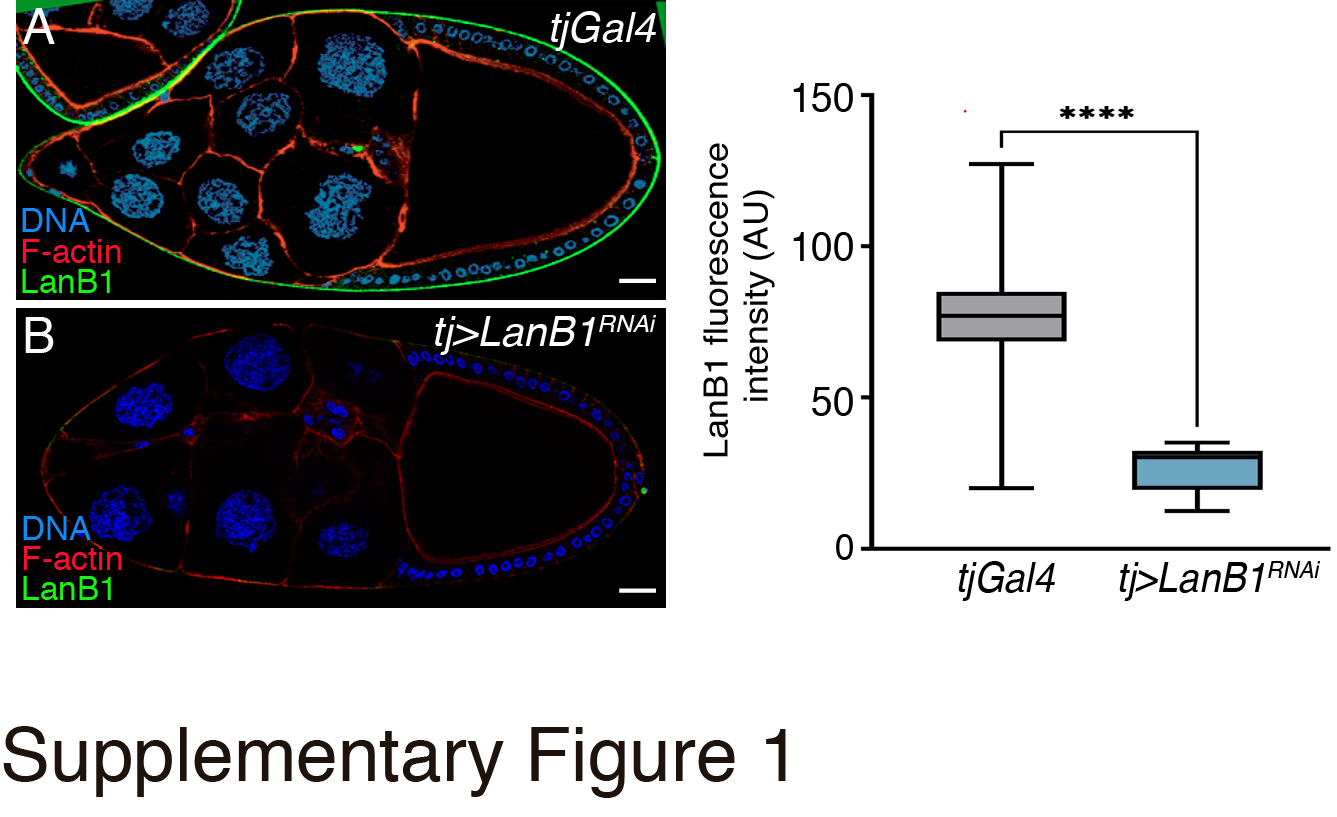

### Sup. Fig.2.tif

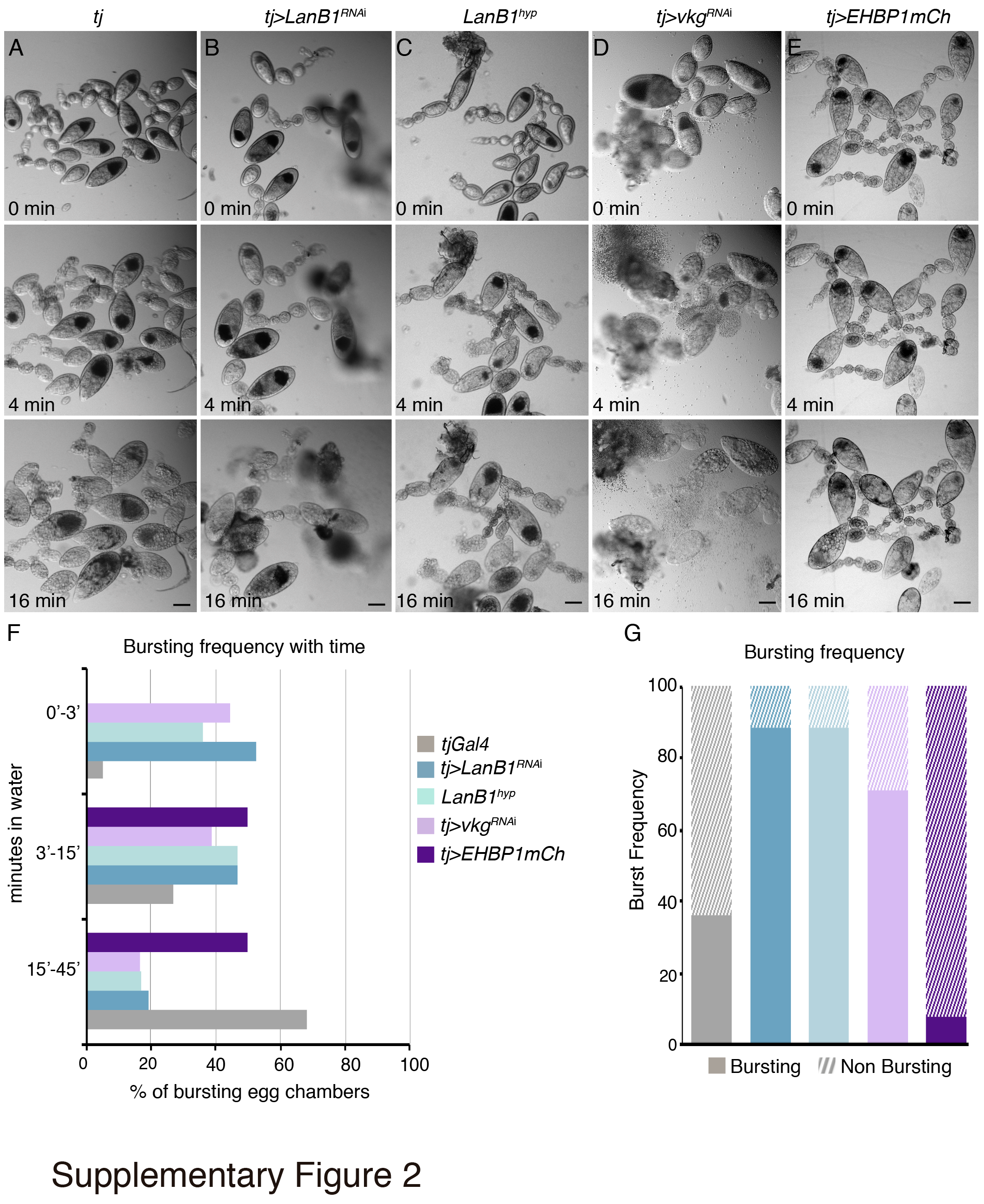

### Sup. Fig.3.tif

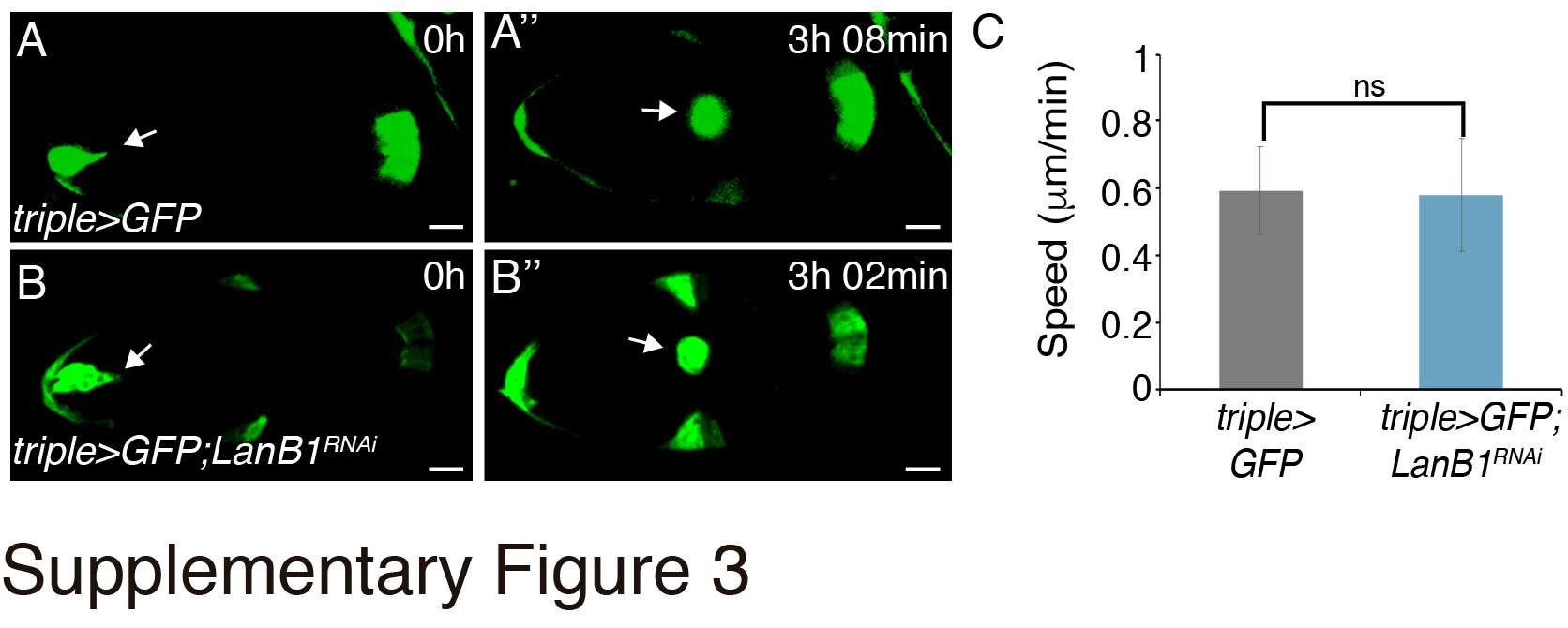

### Sup. Fig.4.tif

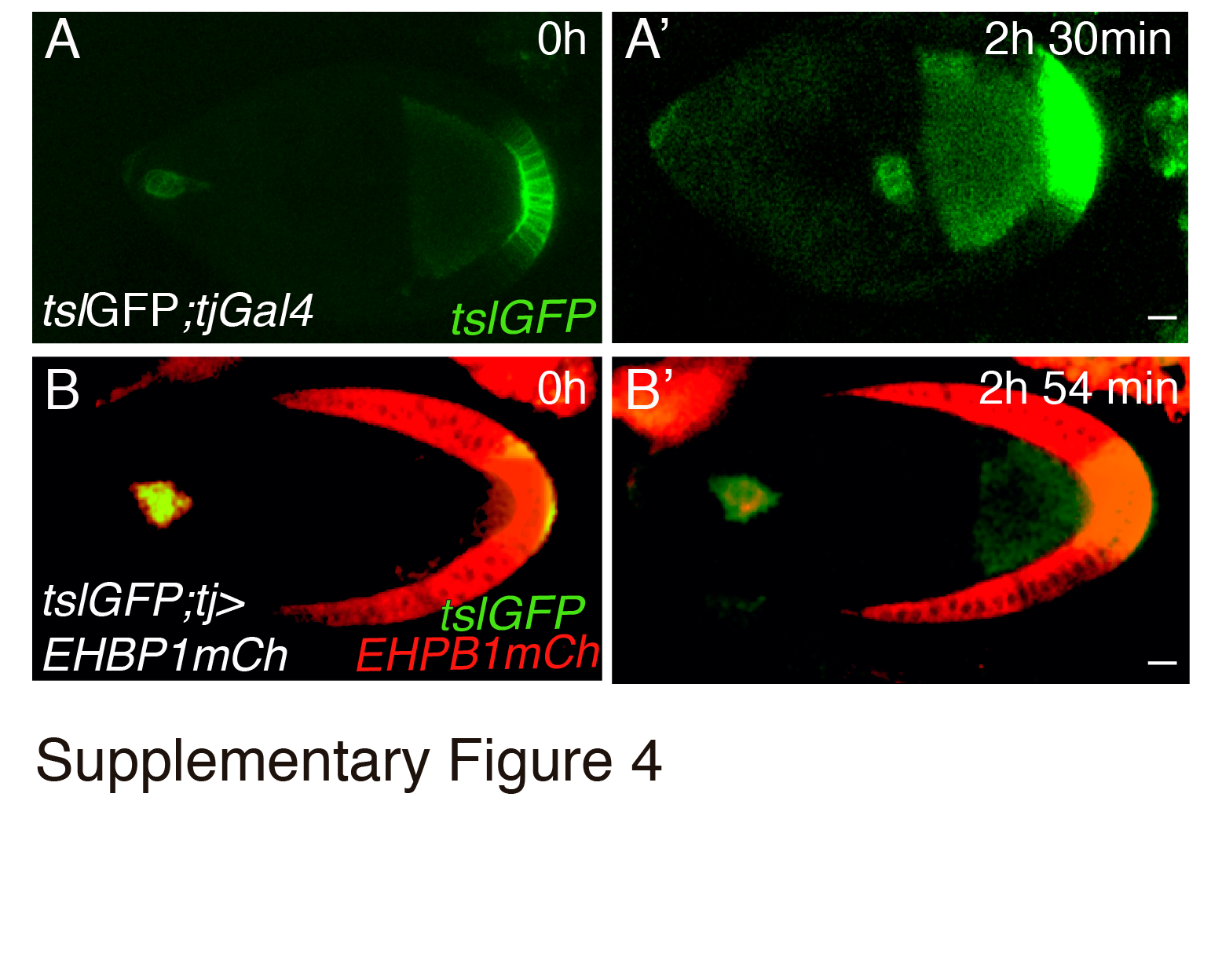

### Sup. Fig.5.tif

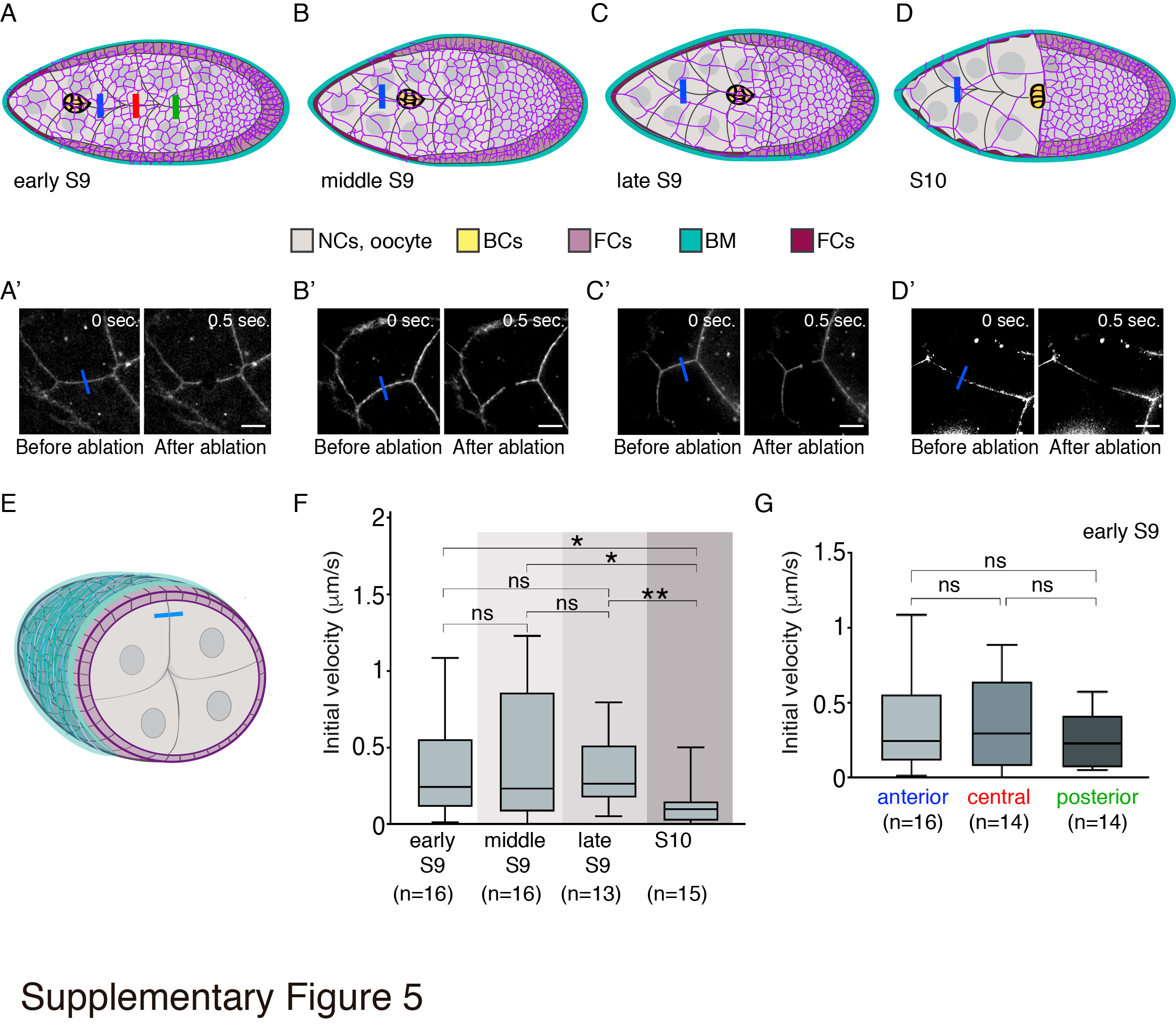

### Sup. Fig.6.tif

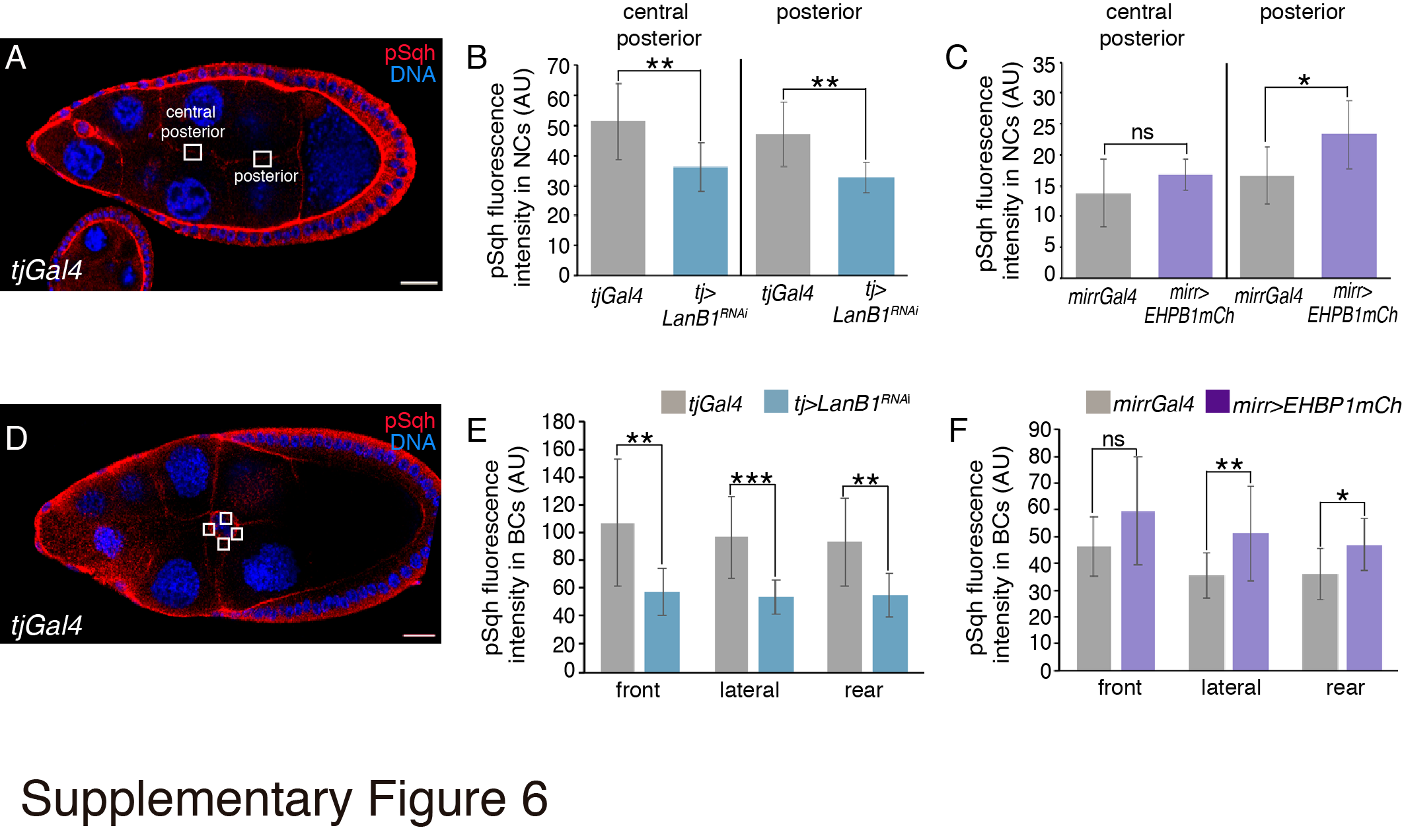

### Sup. Fig.7.tif

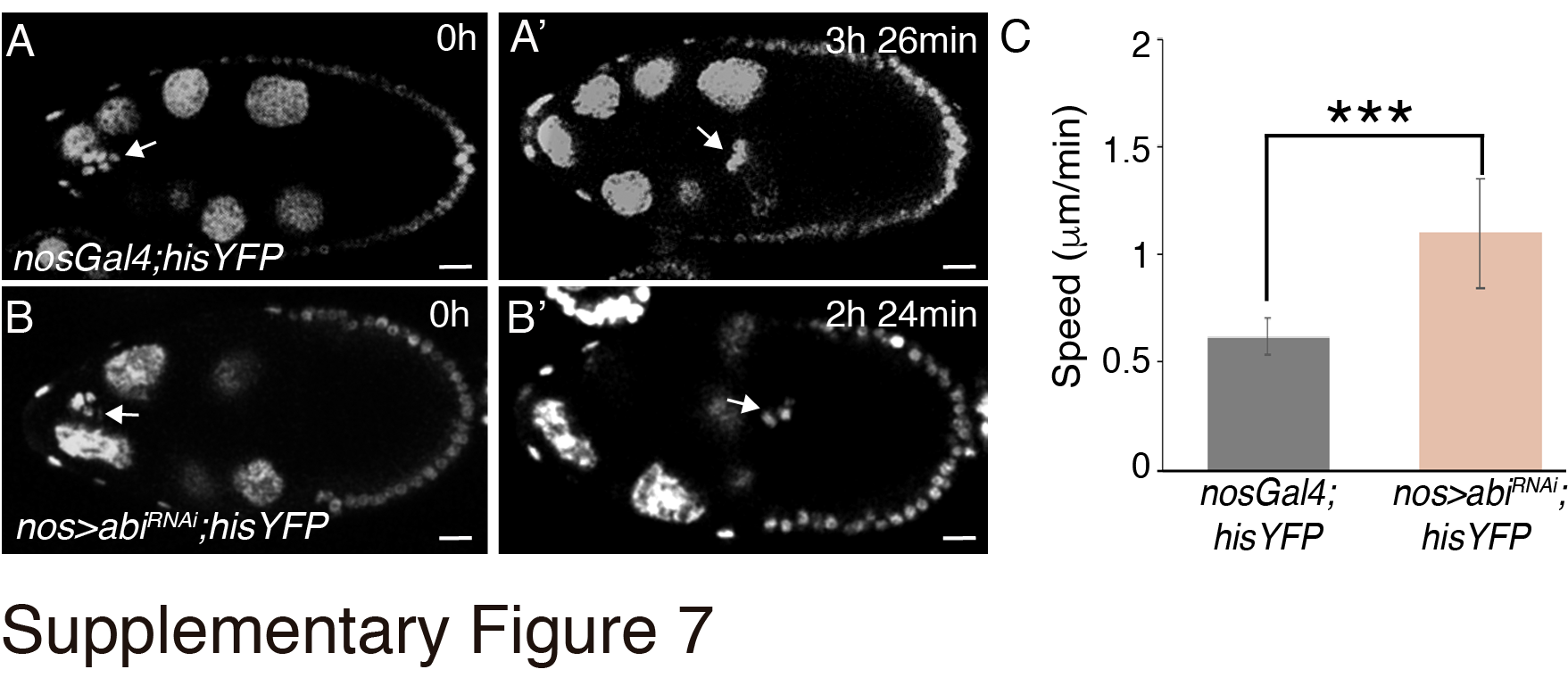

### Sup. Fig.8.tif

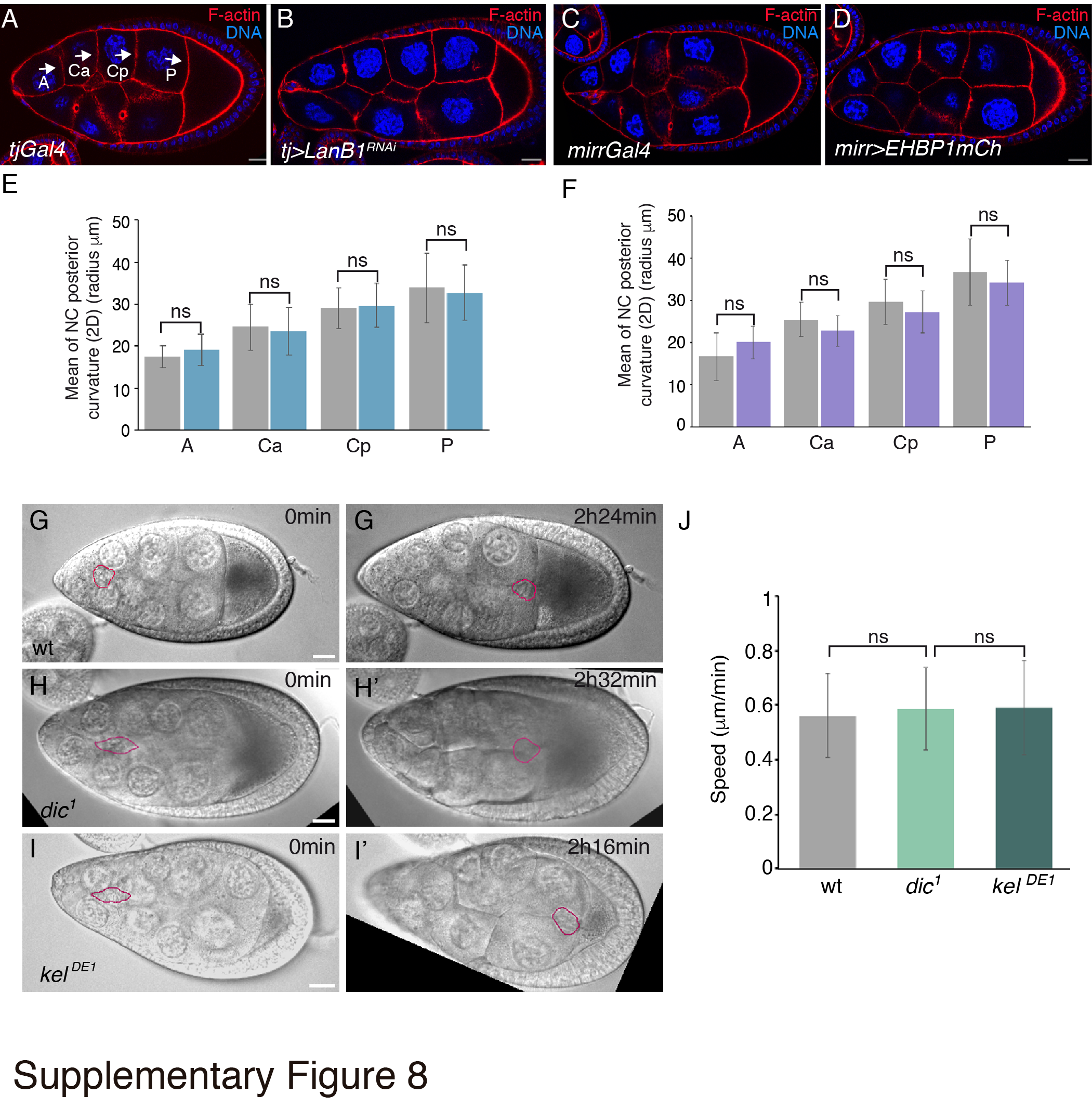

### Sup. Fig.9.tif

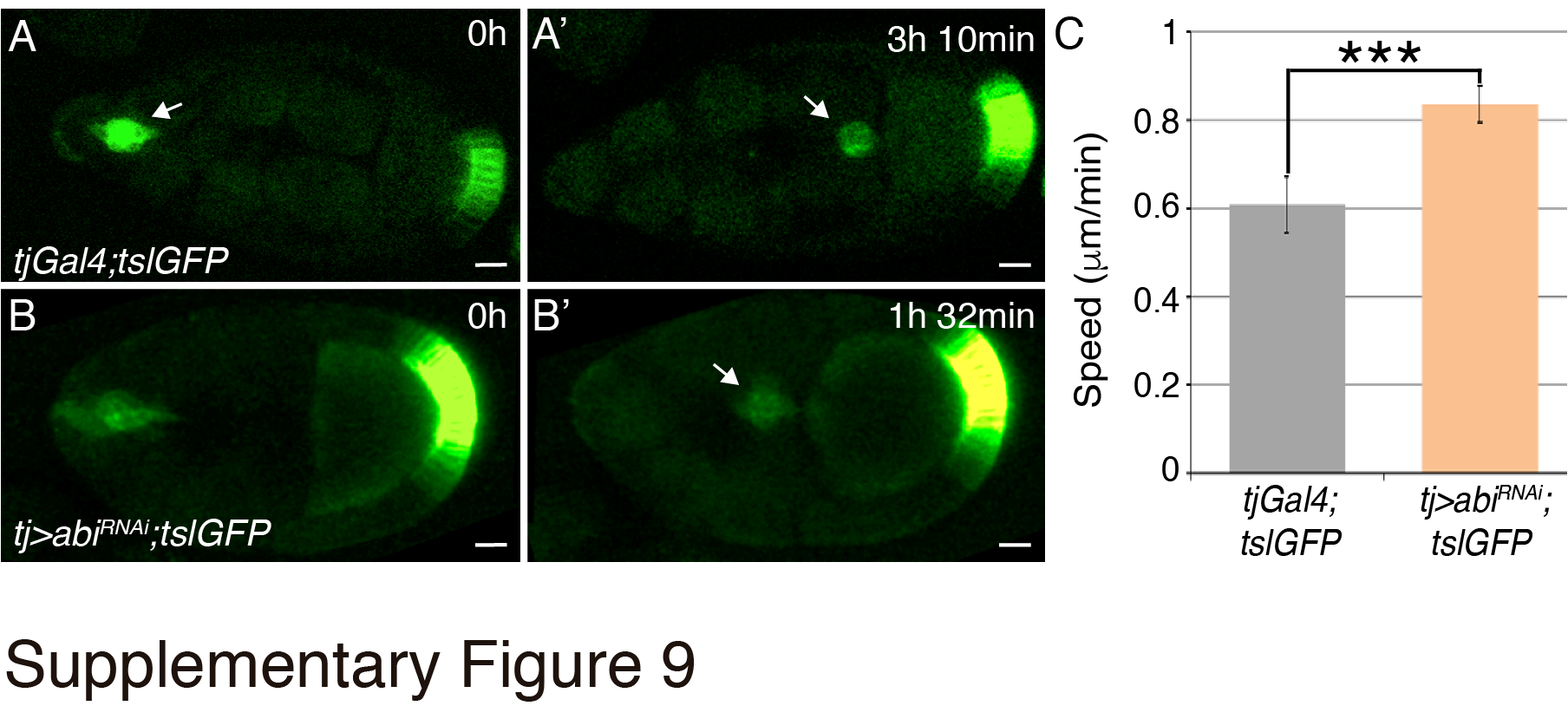
